# Mitofusin-2 in nucleus accumbens D2-MSNs regulates social dominance and neuronal function

**DOI:** 10.1101/2022.10.07.511275

**Authors:** Sriparna Ghosal, Elias Gebara, Eva Ramos-Fernández, Alessandro Chioino, Jocelyn Grosse, Bernard Schneider, Antonio Zorzano, Simone Astori, Carmen Sandi

## Abstract

The nucleus accumbens (NAc) is a brain hub regulating motivated behaviors, including social competitiveness. Mitochondrial function in the NAc is critically implicated in the association between anxiety and social competitiveness, and the mitochondrial fusion protein mitofusin 2 (Mfn2) in NAc neurons has been shown to regulate anxiety-related behaviors. However, it remains unexplored whether accumbal Mfn2 levels also affect social behavior and whether Mfn2 actions in the emotional and social domain are driven by distinct cell types. Here, we found that subordinate-prone highly anxious rats show reduced accumbal Mfn2 levels and that Mfn2 overexpression promotes dominant behavior. In mice, selective Mfn2 downregulation in NAc dopamine D2 receptor-expressing medium spiny neurons (D2-MSNs) induced social subordination, accompanied by reduced mitochondrial function and decreased neuronal excitability. Instead, D1-MSN-targeted Mfn2 downregulation affected competitive ability only transiently mainly by increases in anxiety-like behaviors. Our results assign dissociable cell-type specific roles to Mfn2 in the NAc in modulating social dominance and anxiety.

## Introduction

Social hierarchy among conspecifics governs group organization, where dominant individuals have privileged access to important resources such as food, mating partners and territories (van der Kooij and Sandi, 2015; Sapolsky, 2005). Thus, the benefits associated with social rank provide a strong motivation to become dominant. Conversely, social subordination is associated with important disadvantages for health and wellbeing (Gilbert et al., 1998; Marmot, 2006; Sapolsky, 2005). Accordingly, there is an increasing interest in identifying the (neuro)biological determinants of social rank (Dwortz et al., 2022; Lee et al., 2021; Papilloud et al., 2020; Timmer et al., 2011; Wang et al., 2011; Weger et al., 2018; Zhang et al., 2022; Zhou et al., 2017).

Social dominance is typically established through social competitions (Neumann et al., 2010; Sandi and Haller, 2015; So et al., 2015; Zhou et al., 2017), and the nucleus accumbens (NAc) -a critical element of the brain’s motivation system-is one of the key regions critically implicated in determining the outcome of a social competition (Ghosal et al., 2019; Hollis et al., 2015; van der Kooij et al., 2018; Shan et al., 2022). The NAc integrates inputs from regions involved in the regulation of social dominance, including the ventral tegmental area, hippocampus, amygdala, and prefrontal cortex to regulate social behavior (Dwortz et al., 2022; Gunaydin et al., 2014; Russo and Nestler, 2013; Wang et al., 2011). Two major populations of medium spiny neurons (MSNs) comprise around 95% of NAc neurons, are projection neurons and classically segregated as those mainly expressing dopamine D1 receptors (D1-MSNs), dynorphin and substance P, and those expressing dopamine D2 receptors (D2-MSNs) and enkephalin (Francis and Lobo, 2017; Kreitzer, 2009; Lobo et al., 2006). These two populations have been attributed divergent roles (reward- and aversion-related, respectively) across several modalities (Floresco, 2007; Kravitz et al., 2012; Lobo et al., 2010), including stress-induced depression-like behaviors (Francis et al., 2015), social behaviors (Lobo et al., 2013) and aggression (Aleyasin et al., 2018). Accumbal D1-MSNs have been associated with social interactions (Muir et al., 2018) and their atrophy with stress-induced social avoidance (Fox et al., 2020) and depression-like behaviors (Fox et al., 2020; Francis and Lobo, 2017). Conversely, emerging evidence points at a role for D2-MSNs in the attainment of social dominance (Shan et al., 2022; Yamaguchi et al., 2017) and -in line with the importance of behavioral vigor in social contests-on the invigoration of behavioral responses elicited by cues that predict rewards (Soares-Cunha et al., 2022). Although contextual conditions may influence rank order in social hierarchies (Cordero and Sandi, 2007; Knight and Mehta, 2017), there are also considerable differences in the predisposition of individuals to attain or strive for dominance (Ellyson and Dovidio, 1985; Hollis et al., 2015; Johnson et al., 2012). Emerging evidence points at mitochondrial function in the NAc as a critical neurobiological underpinning controlling rank attainment (Hollis et al., 2015; van der Kooij et al., 2018). Recently, the differential expression of mitofusin 2 (Mfn2) -a mitochondria outer membrane GTPase-in accumbal MSNs was causally implicated in individual differences in anxiety-like and depression-like behaviors and associated differences in accumbal MSN mitochondrial and neuronal structure and function (Gebara et al., 2021). Mfn2 is known to play a major role in sustaining mitochondrial metabolism and energy homeostasis (Filadi et al., 2018) as well as contributing to mitochondria-endoplasmic reticulum connections (Sebastián et al., 2012). However, whether accumbal Mfn2 is causally implicated in rank attainment -i.e., dominant or subordinate-in dyadic encounters, and whether its impact takes place in an MSN cell type-specific manner, is not known. Here, we hypothesized a role for Mfn2 in D2-MSNs and performed studies in both rats and mice. We show that transcript levels of Mfn2 in the NAc positively correlate with social dominance, and neuronal overexpression of Mfn2 in the NAc neurons increases social dominance in submissive-prone animals. Furthermore, we identify a key role for Mfn2 in D2-MSNs, but not D1-MSNs, in social rank attainment, and illustrate a drastic impact of Mfn2 content for D2-MSN mitochondrial and neuronal structure and function.

## Results

### Mfn2 is reduced in NAc D1- and D2-MSNs of subordinate-prone rats, and its overexpression increases social dominance

Figure 1A summarizes the main findings from our previous work in rats in which we identified a crucial role of NAc mitochondrial function in regulating who wins and who loses a social competition between two unfamiliar males, and pinpointed Mfn2 as a key molecule defining mitochondrial, cellular and behavioral phenotypes. Specifically, males prone to become subordinate during a social encounter -identified a priori as those showing a basal predisposition for high anxiety-like (HA) levels-exhibit lower mitochondrial function -manifested by a reduced respiratory capacity, decreased ATP content and increased levels of reactive oxygen species-in the NAc than those -with a basal predisposition for low anxiety-like (LA) levels-prone to become dominant (Gebara et al., 2021; Hollis et al., 2015; van der Kooij et al., 2018). Interfering with accumbal mitochondrial function bidirectionally regulates who wins a social contest. In addition, subordinate-prone rats display as well lower spine density and dendritic arborization in accumbal MSNs and lower levels of mitofusin 2 (Mfn2) in both D1-and D2-MSNs than dominant-prone counterparts (Gebara et al., 2021). This lower Mfn2 content in MSNs in subordinate-prone animals was causally implicated in their enhanced anxiety-like and depression-like behaviors, as well as in the identified deficits in mitochondrial and neuronal structure and function (Gebara et al., 2021).

**Figure 1.**
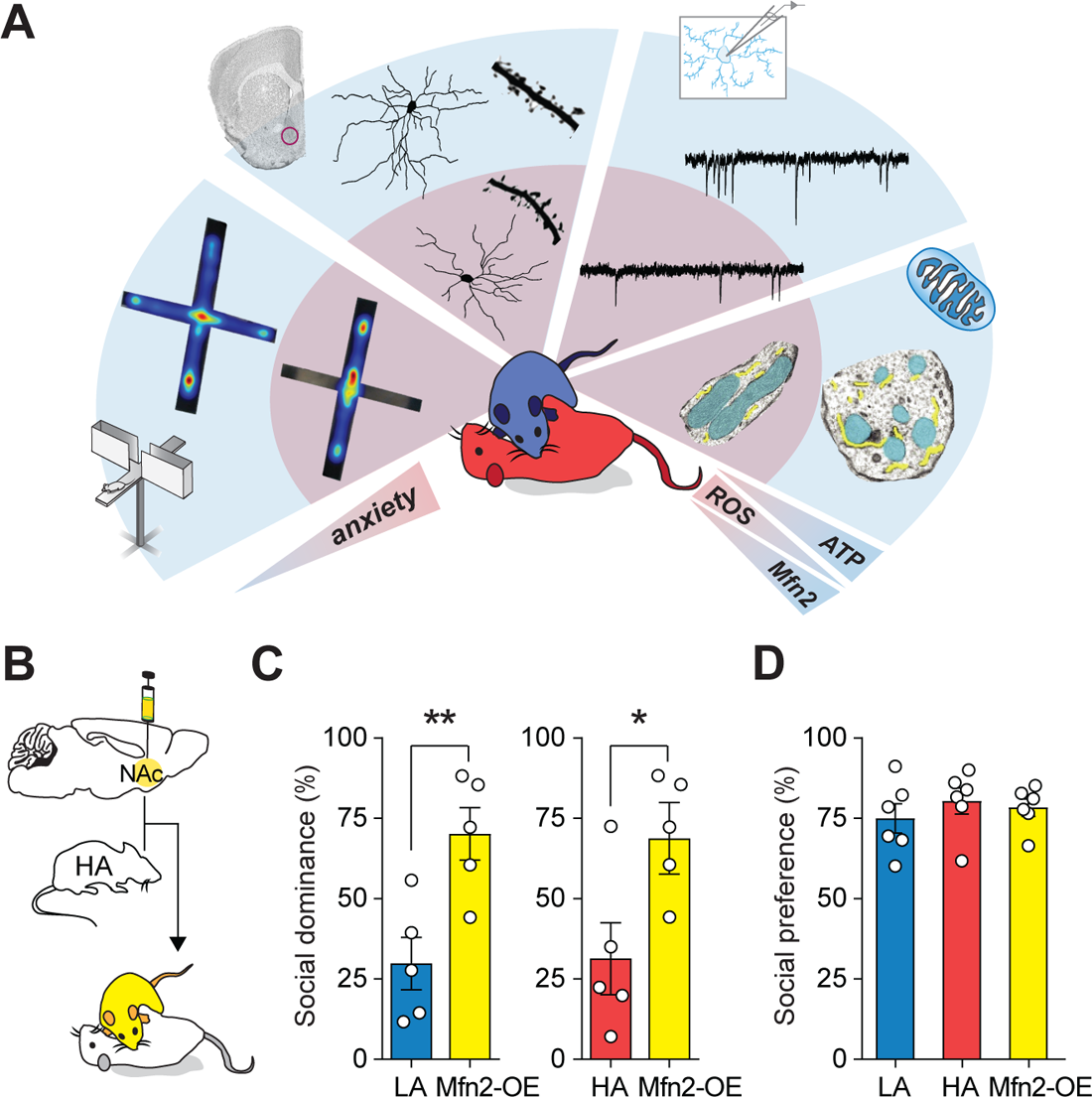
Mfn2 overexpression in the NAc neurons enhances social dominance. (A) Summary of the main findings from social dominance studies in Wistar rats (Gebara et al., 2021; Hollis et al., 2015; van der Kooij et al., 2018). Males prone to lose a social competition (color-coded in red) -identified a priori as highly anxious (HA) based on EPM performance (example EPM heat maps on the left)-displayed a reduction in dendritic arborization, spine density and excitatory synaptic inputs in NAc MSNs, as compared to dominant low anxious (LA) males (color-coded in blue). In the NAc, HA rats presented mitochondria (cyan) with elongated globular shape, fewer contacts with the endoplasmic reticulum (yellow), accompanied by reduced respiratory capacity, decreased ATP content and increased levels of reactive oxygen species (ROS). Transcript levels of Mfn2 were significantly lower in NAc MSNs from HA rats compared to LA rats. (B) HA rats were injected with a Mfn2-overexpressing AAV into the NAc (Mfn2-OE) followed by confrontation with sham-operated HA or LA rats. (C) Mfn2-OE rats showed increased social dominance when confronted with sham-operated LA and HA rats. (D) There were no differences in social preference regardless of Mfn2 overexpression or trait anxiety. Data are presented as mean ± SEM. * p < 0.05, ** p < 0.01. For detailed statistical information, see the Table S1. This figure is complemented by Figure S1.

Here, we investigated the role of accumbal Mfn2 in social dominance. We classified male Wistar rats as LA or HA according to their behaviors in anxiety-related tasks (Figures S1A-D). Consistent with our previous results (Hollis et al., 2015), HA rats exhibited reduced offensive behavior during the social encounter (Figure S1E). Importantly, there was a significant positive correlation between social dominance level and the expression of Mfn2 in the NAc (Figure S1F). Similarly, social dominance level positively correlated with MSN dendritic complexity (Figure S1F).

To ascertaining whether low NAc Mfn2 levels in subordinate-prone/HA animals are at the core of their disadvantage to win a social competition, we virally induced Mfn2 overexpression (Mfn2-OE) by injecting a pAAV9-hSyn1-Myc-hMFN2 vector (Gebara et al., 2021) into the NAc of HA rats (Figure 1B). Mfn2-OE HA rats showed higher social dominance levels in a confrontation with another sham-operated LA or HA rat, indicating that restoring Mfn2 in NAc neurons reverses their predisposition to lose a social contest (Figure 1C). To examine whether this difference in social dominance was related to social anxiety, we tested LA, HA and OE rats in a social preference paradigm and found no group differences in the time spent exploring a juvenile rat over an inanimate object (Figure 1D). Collectively, these results establish that the odds of winning a social competition are regulated by Mfn2 content in NAc neurons.

### Mfn2 downregulation in the NAc D2-MSNs leads to social subordination

To address the question of whether the Mfn2 level in D2-MSNs plays a role in social dominance, we injected a Lenti virus expressing Cre recombinase under the enkephalin (Enk) promoter (Zhang et al., 2015) or a Lenti-Enk-GFP (GFP) into the NAc of Mfn2 floxed (Mfn2^f/f^) mice (Chen et al., 2007) (Figure 2A). Enk-Cre injected mice (termed as Enk^Cre^) showed a reduction of *Mfn2* expression in the NAc (Figure 2B). Enk^Cre^ and GFP mice were matched for body weight and placed together to cohabitate in a cage in dyads, including one animal from each group (Enk^Cre^ and GFP). Two weeks after the beginning of their cohabitation, they underwent the social confrontation tube test (Figure 2C). The tube test results demonstrate that Mfn2 downregulation in D2-MSNs reduced social rank, as revealed by the total number of wins across all sessions and in each trial (Figures 2D and 2E). Notably, the social rank assessed in the tube test in the Enk^Cre^ was not due to any learning deficits during the training phase as both Enk^Cre^ and GFP mice spent similar amount of time in the tube during the training phase (Figure S2A), or to any effect of Mfn2 knockdown on muscle strength (Figure S2B) or body weight (Figure S2C). Moreover, plasma corticosterone (Figure S2D) and testosterone (Figure S2E) levels were similar between the two groups. Importantly, further analysis of the mice’s actions in the tube test indicated that the lower rank of the Enk^Cre^ mice was attributable to a decrease in the relative time spent on the push action (Figures 2F and 2G) and to an increase in the time spent on retreat actions (Figures 2H and 2I). To further evaluate the role of Mfn2 in D2-MSNs in social dominance, we performed the warm spot test (Figure 2J), which models ethological competition for important resources. The Enk^Cre^ mice demonstrated a reduction in the relative time occupying the warm spot (Figure 2K).

**Figure 2.**
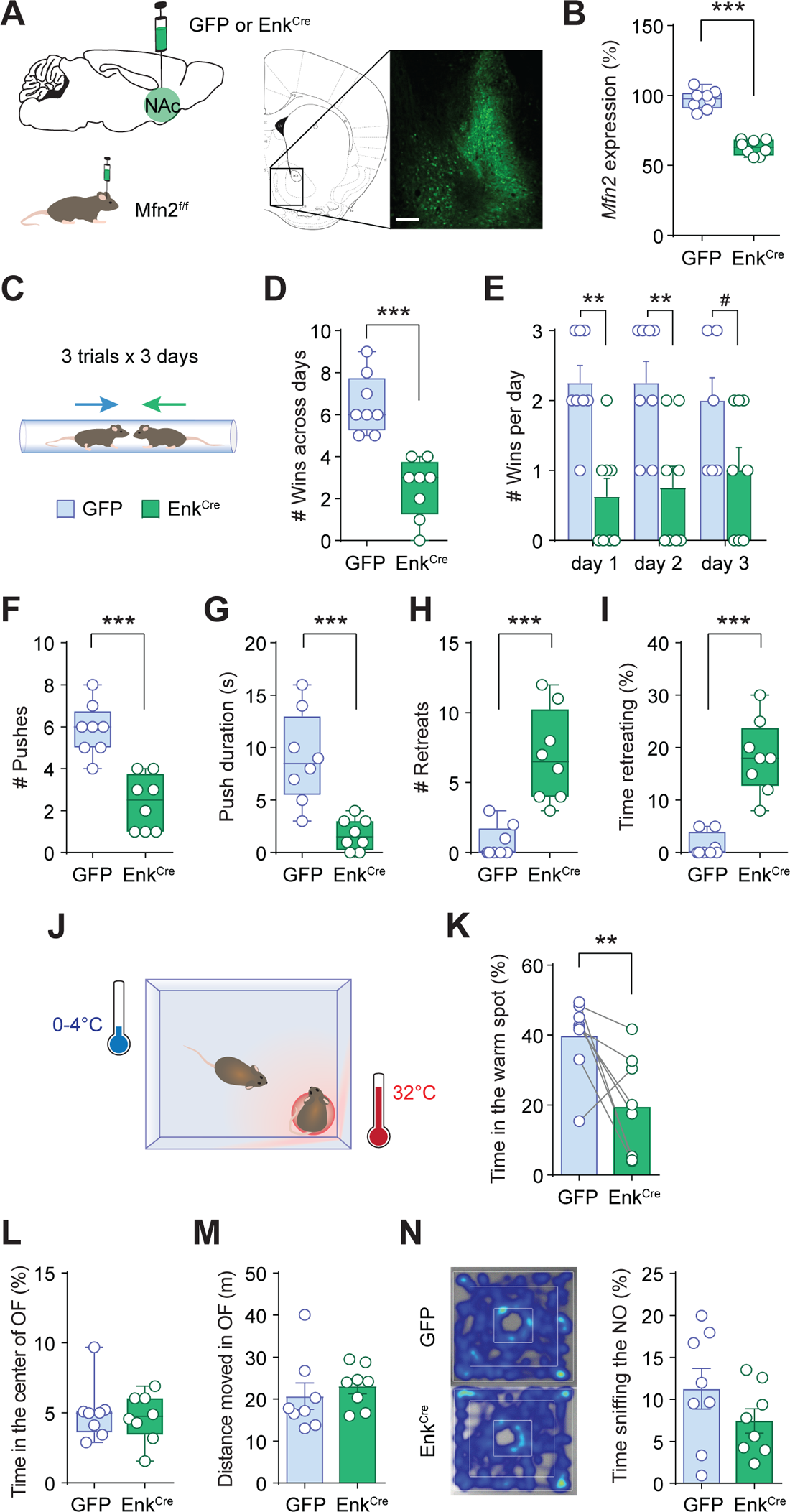
Mfn2 knockdown in NAc D2-MSNs promotes social subordination. (A) Mfn2^f/f^ mice received bilateral infusion of Lenti-Enk-GFP or Lenti-Enk-Cre into the NAc. Right: representative image of GFP fluorescence showing targeted NAc region (scale bar = 100 µm). (B) *Mfn2* expression levels in the NAc indicate efficient Mfn2 knockdown in Enk^Cre^ mice. (C) Schematic showing experimental approach for the social confrontation tube test. (D) The number of wins was significantly decreased in the Enk^Cre^ across all the tube test trials. (E) Across all sessions in the tube test, Enk^Cre^ mice showed lower winning behavior. (F-G) Both the frequency of pushes as well as the average duration of pushing behavior were significantly reduced in the Enk^Cre^ mice. (H-I) Both the frequency of retreats as well as the % duration retreating were increased in the Enk^Cre^ mice. (J) Schematic showing experimental approach for the warm spot test. (K) Enk^Cre^ mice spent less time in the warm spot. (L-M) Time spent in the center of the OF and total locomotion in the OF showed no difference between GFP and Enk^Cre^ mice. (N) The time spent in the center interacting with the novel object (NO) showed no difference between GFP and Enk^Cre^ mice. Left: Representative heatmaps of mouse position during the NO test. Data are mean ± SEM in bar graphs or min-to-max in box-and-whisker plots. Circles represent single observations (n = 8/ group). # p < 0.1, ** p < 0.01, *** p < 0.001. Exact statistics can be found in Table S1. This figure is complemented by Figure S2.

Given our previous findings linking anxiety and social dominance (Hollis et al., 2015; van der Kooij et al., 2018), we explored whether Mfn2 downregulation in the D2-MSNs influenced anxiety-like behaviors. No significant differences were observed in the time spent in the center of the open field (Figure 2L), locomotor activity in the open field (Figure 2M) or time spent sniffing a novel object (Figure 2N). Collectively, these data indicate that Mfn2 in the NAc D2-MSNs is necessary for the establishment of social dominance, but does not affect trait anxiety.

### Mfn2 downregulation in NAc D1-MSNs does not affect social rank

We next examined the functional consequences of NAc D1-MSN specific downregulation of Mfn2. We injected an AAV expressing Cre recombinase under the dynorphin (Dyn) promoter (Dyn^Cre^) (Darvas and Palmiter, 2015) or an AAV-synapsin-GFP (GFP) into the NAc of Mfn2^f/f^ mice (Figure 3A). Bilateral infusion of Dyn^Cre^ caused almost a 50% loss of *Mfn2* expression in the NAc (Figure 3B). Next, to determine whether Mfn2 in D1 causally regulates social dominance, Dyn^Cre^ mice were pair-housed with GFP mice of similar weight. During the training phase, both Dyn^Cre^ and the GFP mice learned to walk through the tube at an equal rate (Figure S3A), ruling out the possible effects of Mfn2 knockdown on learning the task as a confounding factor. Following two weeks of cohabitation, the social hierarchy was tested in the tube test. Across all sessions, GFP and Dyn^Cre^ mice exhibited an equal probability to become dominant and an equal victory rate (Figure 3C), though, there was a lower victory rate in the Dyn^Cre^ mice in the first encounter, as evidenced by fewer number of wins only in the first confrontation (Figure 3D). To further evaluate the role of Mfn2 in D1-MSNs in social dominance, we performed the warm spot test. We found both Dyn^Cre^ and GFP mice spent similar amount of time in the warm spot (Figure 3E). No differences were found in the body weight (Figure S3B) or muscle strength (Figure S3C), plasma corticosterone (Figure S3D) or plasma testosterone (Figure S3E) levels between the Dyn^Cre^ and GFP mice, ruling out the possible effect of size, corticosterone, or androgen as a confounding factor on the observed behavioral effects of Mfn2 downregulation in the NAc D1-MSNs. These data indicate Mfn2 in the NAc D1-MSNs is necessary for the establishment of social dominance only in the first confrontation, but does not affect the stabilization of social ranking.

**Figure 3.**
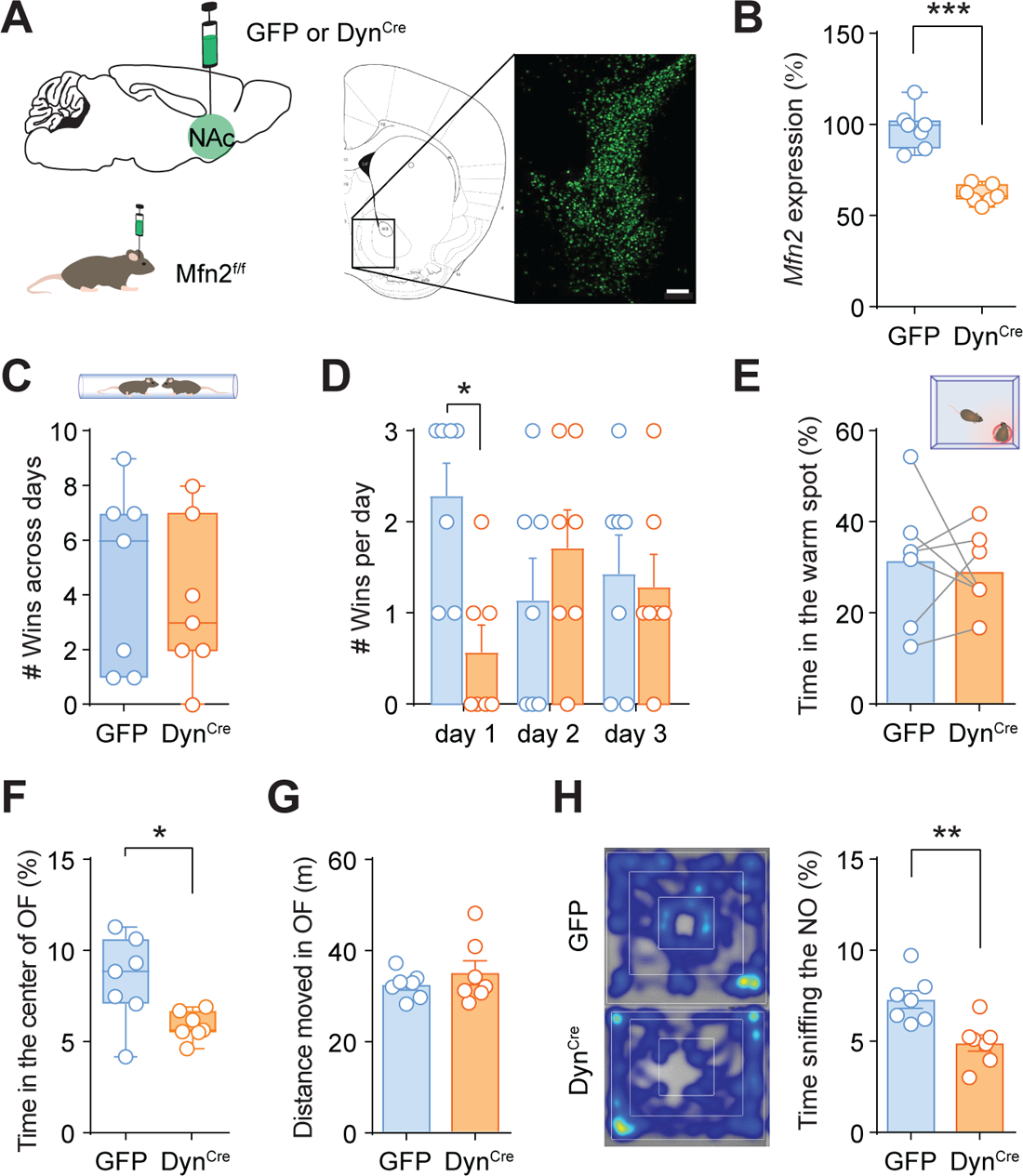
Mfn2 knockdown in NAc D1-MSNs does not alter social rank but promotes anxiety-like behaviors. (A) Mfn2^f/f^ mice received bilateral infusion of AAV-Syn-GFP or AAV-pDyn-Cre in the NAc. Right: representative image of GFP fluorescence shows targeted NAc region (scale bar = 100 µm). (B) *Mfn2* expression levels in the NAc indicate efficient Mfn2 knockdown in Dyn^Cre^ mice. (C) Across all sessions of tube test, there were no differences between GFP and Dyn^Cre^ mice in the number of trials won. (D) Dyn^Cre^ mice showed fewer wins on the first day of tube test competition. (E) There was no difference in the time spent in warm spot between GFP and Dyn^Cre^ mice. (F) Dyn^Cre^ mice showed enhanced anxiety-like behaviors, as measured by reduced time spent in the center of the OF. (G) Total locomotion was not affected in Dyn^Cre^ mice. (H) NO tests showed reduced NO exploration in the Dyn^Cre^ mice. Left: representative heatmaps of mouse position during the NO test. Data are mean ± SEM in bar graphs or min-to-max in box-and-whisker plots. Circles represent single observations (n= 7/ group). * p < 0.05, ** p < 0.01, *** p < 0.001. Exact statistics can be found in Table S1. This figure is complemented by Figure S3.

The mice were then characterized for anxiety and exploratory behaviors. Compared to GFP mice, Dyn^Cre^ mice spent less time in the center of an open field (Figure 3F) and less time sniffing a novel object in the novel object reactivity test (Figure 3H), which was not caused by changes in locomotor activity (Figure 3G).

Altogether, our results indicate that Mfn2 in NAc D1-MSNs has minimal impact on the attainment and expression of social dominance, however, it is necessary for the expression of anxiety-like behaviors.

### Mfn2 downregulation alters mitochondrial function in D2-MSNs

Given our previous findings linking bioenergetics in the NAc with the differential predisposition to win a social competition (Hollis et al., 2015; van der Kooij et al., 2018), we explored whether Mfn2 downregulation in D2-MSNs affected NAc mitochondrial respiration (Figure 4A). Enk^Cre^ mice showed significantly lower mitochondrial respiration through complexes I and II compared to GFP controls (Figure 4B). Furthermore, as the major output of mitochondrial function is ATP production, we measured total levels of ATP in the NAc tissue homogenates. Although no significant differences were present in the ATP concentration or the total amount of nucleotide (ATP+ADP) between Enk^Cre^ mice and the GFP controls, we observed a significant reduction in the ADP concentration and the ATP/ADP ratio in the Enk^Cre^ mice (Figures 4C-E), suggesting an impaired mitochondrial oxidative phosphorylation in the NAc of Enk^Cre^ mice. Previously, we highlighted the necessity of Mfn2 in maintaining the contact between mitochondria and ER in the MSNs (Gebara et al., 2021). Thus, we examined ER–mitochondria interactions within the NAc in Enk^Cre^ and GFP mice using proximity ligation assay (PLA), that quantifies the interaction of the mitochondrial VDAC1 protein and the ER protein IP3R (Figure 4F). Immunofluorescence staining illustrates the distribution of labeled VDAC1-IP3R interacting dots, reflecting ER–mitochondria contacts (Figure 4F), which were significantly decreased in Enk^Cre^ mice (Figure 4G).

**Figure 4.**
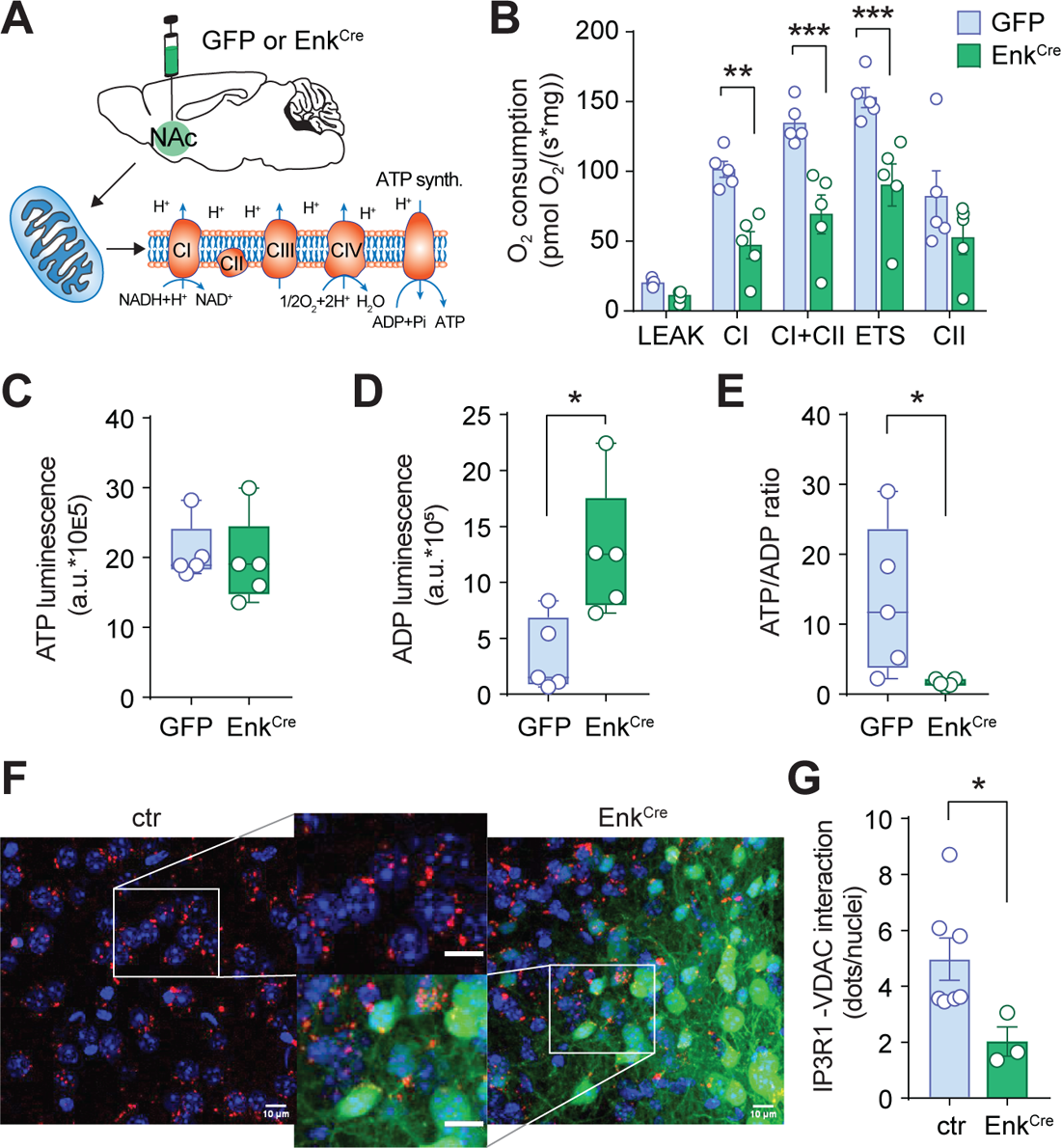
Mfn2 knockdown in NAc D2-MSNs decreases overall mitochondrial function. (A) Schematic showing assessment of mitochondrial respiratory chain function in GFP and Enk^Cre^ mice. (B) Enk^Cre^ mice showed reduced mitochondrial respiration in the NAc. (C-E) Levels of ATP and ADP and ATP/ADP ratio in the NAc of GFP and Enk^Cre^ mice. (F) Representative fluorescence images showing VDAC1-IP3R1 interacting dots (red), DAPI staining (blue), Cre positive cells (green) in NAc from Mfn2^f/f^ (control, Ctr) and Enk^Cre^ mice, and insets with higher magnification (scale bars = 10 μm). (G) Quantification of VDAC1-IP3R1 interacting dots, measured over Cre positive and Ctr cells. Data are mean ± SEM in bar graphs or min-to-max in box-and-whisker plots. Circles in the bar graphs represent single observations. * p < 0.05, ** p < 0.01, *** p < 0.001. Exact statistics can be found in Table S1. This figure is complemented by Figure S4.

Recent studies have reported hippocampal and cortical neuronal degeneration with aging in Mfn2 knockout mice compared with age-matched littermate control mice (Han et al., 2020; Jiang et al., 2018). We, therefore, examined gross morphology of the NAc region and neuronal viability in the Enk^Cre^ and Dyn^Cre^ mice used in the behavioral assays. Mfn2 knockdown did not affect NAc morphology, as shown by Hematoxylin-eosin (HE)-staining (Figures S4A-C). Moreover, NeuN immunoreactivity was also intact (Figures S4D and S4D’), demonstrating that the behavioral phenotype is specific to Mfn2 knockdown, and unrelated to any gross neuronal damage. Furthermore, Mfn2 knockdown in the NAc MSNs did not precipitate cell death in these regions beyond that of a control level, as evidenced by cleaved caspase 3 immunoreactivity (Figures S4E and S4E’). Collectively, these data indicate that delivery of the respective Cre-expressing viruses can effectively reduce *Mfn2* expression levels in the NAc region without affecting neuronal viability.

### Mfn2 downregulation causes reductions in dendritic complexity and excitability in D2-MSNs

To explore whether differences in social dominance are associated with structural and functional differences in MSNs of the NAc, as previously shown for the HA rats (Gebara et al., 2021), we performed morphometric analysis of Golgi-impregnated neurons in mice with Mfn2 deletion selectively in the D2 cells (Mfn2^Adora2a^). MSNs in Mfn2^Adora2a^ mice displayed a marked dendritic atrophy, as demonstrated by decreased Sholl intersection profile (Figure 5B) and reduced total dendritic length (Figure 5C).

**Figure 5.**
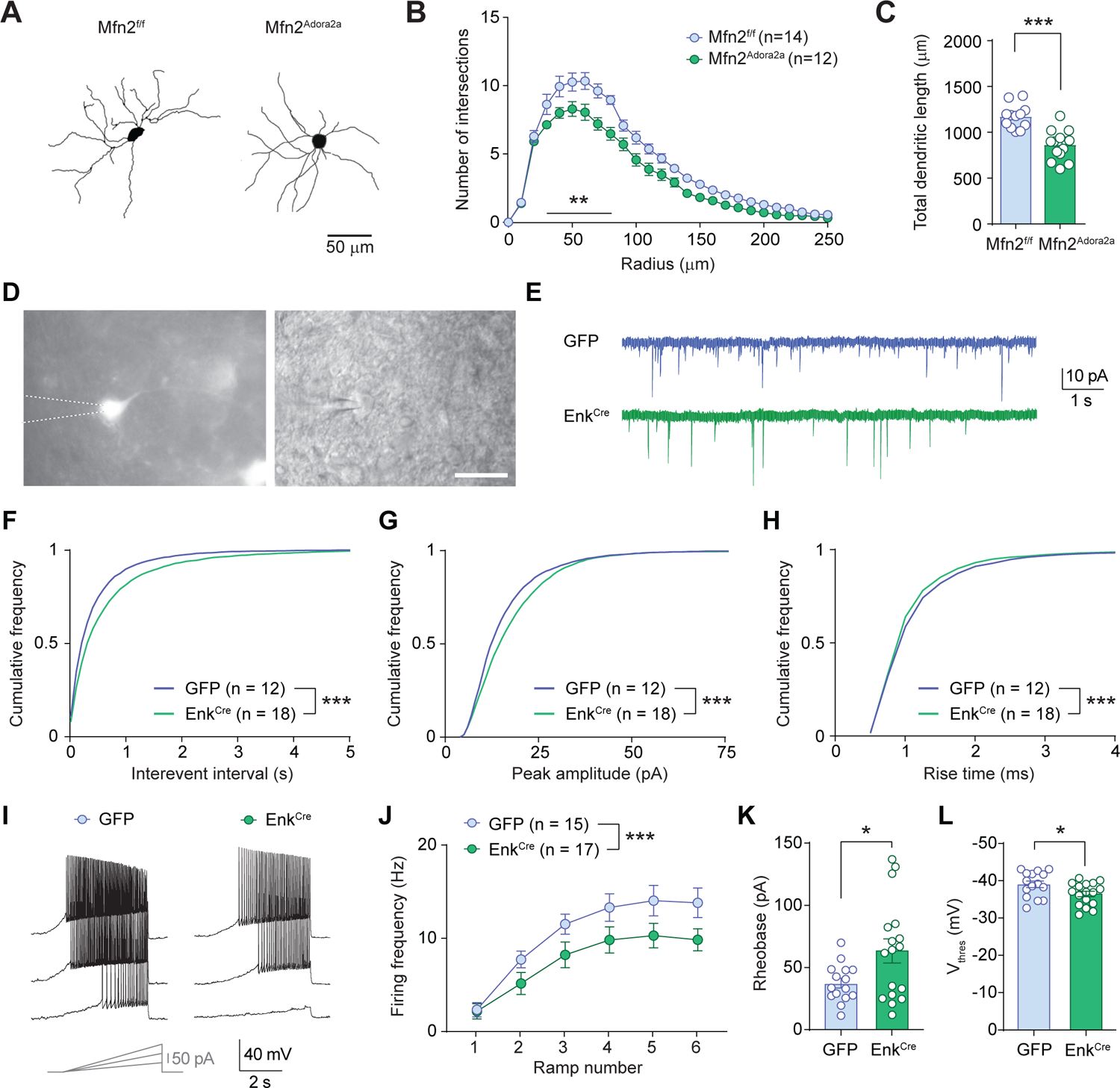
Mfn2 knockdown in NAc D2-MSNs decreases dendritic complexity and neuronal excitability. (A) Reconstructions of Golgi-stained MSNs in NAc from Mfn2^f/f^ and Mfn2^Adora2a^ mice. (B-C) Sholl intersection profile indicates reduced dendritic complexity, accompanied by decreased total dendritic length of MSNs in Mfn2^Adora2a^ compared with Mfn2^f/f^ mice. (D) Micrographs showing GFP fluorescence and transmitted light images of D2-MSNs recorded with a patch pipette in acute slices from GFP and Enk^Cre^ mice (scale bar = 20 µm). (E) Example traces showing miniature excitatory postsynaptic currents (mEPSCs) recorded in infected D2-MSNs from GFP and Enk^Cre^ mice. (F-H) Cumulative distribution plots interevent interval, peak amplitude and rise time of mEPSCs recorded in infected D2-MSNs from GFP and Enk^Cre^ mice. (I-J) Example traces showing D2-MSN firing elicited by somatic injection of current ramps. Enk^Cre^ mice displayed a significantly reduced frequency of discharge compared to GFP mice. (K-L) Minimal current injected to elicit the first action potential (rheobase) and firing threshold in D2-MSNs from GFP and Enk^Cre^ mice. Data are mean ± SEM, and circles in the bar graphs represent single observations for panel C. For panels F-J, statistical difference was tested with the Kolmogorov-Smirnov test. * p < 0.05, ** p < 0.01, *** p < 0.001. Exact statistics can be found in Table S1.

To further explore the functional consequences of these morphological changes for D2-MSNs excitability, we performed *ex vivo* patch-clamp recordings from infected D2-MSNs in acute slices from GFP and Enk^Cre^ mice (Figure 5D). Miniature excitatory postsynaptic currents (mEPSCs) were less frequent in Enk^Cre^ mice, as indicated by the overall larger interevent interval values (Figure 5E), suggesting a reduction in the number of glutamatergic synaptic inputs. The distribution of the peak amplitude values of mEPSCs displayed a shift towards larger values in Enk^Cre^ mice (Figures 5F and 5G). This effect may be due to the fact that proximal inputs of apparent larger amplitude (as measured in somatic patch-clamp recordings) contribute more to the population of events acquired in Enk^Cre^ D2-MSNs, as opposed to distal inputs, which are expected to be less represented in MSNs with reduced dendritic complexity. To assess this possibility, we measured mEPSCs rise time and found that, indeed, it was faster in D2-MSNs from Enk^Cre^ mice than in GFP (Figure 5H), indicating that proximal inputs (with kinetics that are less affected by cable filtering properties) are more dominant in Enk^Cre^ mice. When neuronal firing was elicited by depolarizing current injections, D2-MSNs from Enk^Cre^ mice displayed a significantly reduced frequency of discharge (Figures 5I and 5J) and required larger current injections (Figure 5K) to reach the firing threshold, which was more depolarized as compared to GFP mice (Figure 5L). Altogether, these data indicate the Mfn2 downregulation causes a net decrease in D2-MSNs excitability, deriving from both a reduction in the density of excitatory synaptic inputs and a deficit in sustaining neuronal firing.

## Discussion

Our study highlights a key role for the mitochondrial GTPase Mfn2 in NAc D2-MSNs in the regulation of social dominance. Using a model of natural phenotypic variation in outbred rats, we causally implicate a lower accumbal Mfn2 neuronal content in HA animals in their disadvantage to win a social competition against less anxious ones. Then, by applying a cell-type specific downregulation of Mfn2 in mice, we identify D2-, but not D1-, MSNs as the neuronal type in which a Mfn2 deficit leads to social subordination. We also show that Mfn2 downregulation in D2-MSNs recapitulates mitochondrial and neuronal alterations observed in the NAc of high anxious rats.

We had previously reported that interindividual differences in anxiety-like behaviors in outbred rats are causally linked to different basal levels of accumbal Mfn2, and that these differences in Mfn2 levels regulate accumbal differences in mitochondrial (i.e., maximal respiratory capacity, volume, and interactions with the ER) and neuronal (i.e., dendritic complexity, spine density, and frequency of excitatory inputs) structure and function in MSNs (Gebara et al., 2021). Here, we show that pan-neuronal Mfn2 overexpression in HA rats drastically enhances their capacity to win a social competition, not only against control HA rats, but also against LA ones which are normally superior over control HA rats. Interestingly, these differences in social competitiveness were not due to differences in social interest or sociability. A key strength of these observations in outbred rats is that they allow explaining the molecular and neurobiological underpinnings of individual differences in a complex behavior in an heterogenous genetic background, as the lack of genetic diversity is a drawback inherent to the use of genetically-homogeneous inbred mice. However, subordinate-prone rats display lower Mfn2 levels in both D1- and D2-MSNs (Gebara et al., 2021), and interrogating the specific cell-type involved in the observed effects requires using the *Cre-LoxP* system for which current tools are optimized in inbred mice. This is particularly relevant, given that NAc D1- and D2-MSN subpopulations are known to divert in their regulation by stress (Fox et al., 2020; Francis and Lobo, 2017), behavioral outcomes (Floresco, 2007) and mitochondrial characteristics (Chandra et al., 2019). Thus, using a *Cre-LoxP* viral-mediated approach in mice, we show a drastic impact of the specific downregulation of Mfn2 in D2-, but not D1-, MSNs on social dominance, as indicated by clear signs of subordination in Enk^Cre^ mice in the tube and the warm spot tests. Importantly, body weight, anxiety-like or locomotor behaviors did not differ between Enk^Cre^ and GFP mice, discarding general body changes to explain the observed differences in social rank. Detailed analyses revealed that the rank degrading induced by Mfn2 knockdown in the D2-MSNs involved both, increased retreats and decreased effortful pushes.

The medial prefrontal cortex (mPFC) has emerged as a key node in the neural circuitry that regulates social competition (Dwortz et al., 2022; Wang et al., 2011), and our results in Enk^Cre^ mice resemble data on the immediate impact of optogenetic inhibition of dorsomedial PFC [dmPFC; including the prelimbic (PL) region and anterior cingulate (ACC) cortex)] pyramidal neurons in their moment-to-moment competitive reactions in the tube test (Wang et al., 2011; Zhang et al., 2022; Zhou et al., 2017). Interestingly, the NAc is a key node in the corticostriatal pathway and receives a great number of afferents from the two main dmPFC subdivisions, the PL and ACC (Gabbott et al., 2005). Prelimbic and ACC projections to the NAc have been critically implicated in the modulation pain perception (Lee et al., 2015; Zhou et al., 2018) and the social transfer of pain (Smith et al., 2021), two processes that are likely to play important roles in the outcome of social competitions and on the maintenance of social hierarchies. Moreover, NAc-projecting PL neurons receive dense inputs from the mediodorsal thalamus (MDT) (Cruz et al., 2021), and inputs from the MDT to the dmPFC have been shown to mediate long-lasting changes in social rank triggered by the accumulated impact of prior winning contests (Zhou et al., 2017). Moreover, NAc efferents are closely connected with the MDT. Indeed, one of the main outputs of NAc D2-MSNs is the ventral pallidum (VP) (KOOB and SWERDLOW, 1988; Leung and Balleine, 2015; Root et al., 2015; Smith et al., 2013) whose inhibitory GABAergic neurons project, in turn, to the MDT (O’Donnell et al., 1997; Root et al., 2015) affecting the processing of reward-related cues (Leung and Balleine, 2015). Thus, NAc D2-MSNs appear to be part of a larger multi-component, cortico-striato-thalamic loop critically regulating social competitiveness and -according to our results here-eventually consolidating individuals’ rank. Our results align well with a recent optogenetic study on NAc D2-MSN-VP projections indicating their role in increasing motivational drive in the presence of reward-predicting cues (Soares-Cunha et al., 2022), as such drive is certainly at the core of defining rank within a hierarchy.

Importantly, the social subordination phenotype in Enk^Cre^ mice was accompanied by alterations in key functional features of mitochondrial and neuronal structure and function. The mitochondrial dysfunctions (i.e., decreased maximal respiratory capacity, lower ATP/ADP ratio and reduced interactions with the ER) and the reductions in dendritic complexity and excitatory inputs mimic alterations previously observed in the NAc of subordinate-prone, HA rats with low constitutive MSN levels of Mfn2 (Gebara et al., 2021). Combined with our previous gain-of-function study -in which Mfn2 overexpression was effective to restore anxiety-related organelle and cellular alterations (Gebara et al., 2021)-the loss-of-function data here reported highlight not only a key role for Mfn2 in D2-MSNs, but also the consequent impoverishment of D2-MSNs neuronal cytoarchitecture and limited recruitment by glutamatergic afferents. D2-MSNs in Enk^Cre^ mice showed reduced frequency of excitatory inputs and decreased firing, indicative of a reduced capacity of D2-MSNs to integrate synaptic input impinging on the NAc and to provide sustained output. These findings align well with a recent study indicating that the postsynaptic strength of excitatory synapses onto D2-MSN in the NAc is positively correlated with social dominance, and that DREADD-induced D2-MSN stimulation or inhibition enhances or reduces social dominance, respectively (Shan et al., 2022). D2 antagonist treatment was also found to attenuate social dominance in highly ranking mice and macaques (Yamaguchi et al., 2017), and D2 receptor expression in the striatum has been found to be increased in dominant rodents and non-human and human primates (Jupp et al., 2016; Martinez et al., 2010; Morgan et al., 2002). Therefore, in line with a proposed role for the specific pattern of neuronal stimulation of each MSN subtype in defining whether they signal reward or aversion (Soares-Cunha et al., 2020), alterations in Mfn2 content may regulate value assignment and effortful engagement, two processes at the core of social hierarchy formation (da Cruz et al., 2018; Lozano-Montes et al., 2019). Importantly, along this line, we recently demonstrated that, in mice, stress during prepuberty that leads to deficits in sociability is associated with decreased expression of Mfn2 in the D2-MSNs in the NAc shell regions (Morató et al., 2022). In addition, several studies involving genetic manipulations of a variety of plasticity-related genes have highlighted the involvement of D2-MSNs in shaping defensive and social behaviors (Chandra et al., 2017; Meirsman et al., 2019).

The lack of effect of Mfn2 downregulation in D1-MSNs on the long-term establishment of the social hierarchy adds to a mixed literature on the role of D1-MSNs and D1 receptors (D1R) in social dominance. Prior work in well-established hierarchies similarly reported that systemic administration of a D1R antagonist had minor effects in social dominance in mice (only increasing it in mice at the middle of the hierarchy) and no effect in macaques (Yamaguchi et al., 2017). However, a recent study in mice found that DREADD-induced NAc D1-MSN stimulation or inhibition increased or decreased social dominance, respectively (Shan et al., 2022). It is important to note that these former studies involved acute and transient manipulations of either D1R or D1-MSN function, and their purpose was to challenge an existing social hierarchy, which is radically different from our study in which we assessed the impact of Mfn2 downregulation on the long-term maintenance of the home cage social hierarchy (i.e., we used the tube test to assess the existing home cage hierarchy not as the immediate readout of an acute manipulation). It is important to differentiate between punctual social competitions [typically discrete and stressful events (Buser et al., 2017; Timmer et al., 2011)] and home cage or long-term social hierarchies (the result of accumulated interactions), as traits important for sustaining dominance may differ from those critical to win punctual competitions. We found that Mfn2 downregulation in D1-MSNs induced only a transient effect on the outcome during the first-day of the tube test competition, but not in following sessions. Similarly, the warm spot test indicated no differences between GFP and Dyn^Cre^ mice. Indeed, one key factor in defining the outcome of social competitions is anxiety (Goette et al., 2015; Hollis et al., 2015), a trait highly responsive to novelty (File, 2001) and stress (Weger et al., 2018). Indeed, the higher anxiety of Dyn^Cre^ mice -as indicated by their behaviors in the open field and novel object tests-may have contributed to the temporary loss of tube test tournaments on day 1. This view aligns with our previous findings in which intra-NAc infusion of a D1R agonist facilitated dominance in an acute confrontation between unfamiliar male rats (van der Kooij et al., 2018), and suggests an involvement of D1-MSNs in the regulation of competitive behavior under acute challenges, but not on the formation of stable home cage hierarchies. Although we do not exclude that stress -and associated mechanisms-triggered by social competitions may imping on factors contributing to long-term hierarchy formation (Cordero and Sandi, 2007; Papilloud et al., 2020), traits important for sustaining social dominance (van der Kooij and Sandi, 2015; Pun et al., 2017) may have a stronger weight on the long term.

Collectively, our results demonstrate that social dominance is regulated by Mfn2 content in NAc D2-MSNs, and indicate the loss of dendritic complexity and reduced D2-MSN excitability as a plausible cellular mechanism linking low accumbal Mfn2 levels with social subordination.

## Acknowledgments

The authors would like to thank Dr. Martin Darvas (University of Washington, USA) for supplying the AAV-Pdyn-Cre vectors and Dr. Fan Wang (Duke University Medical Center, USA) for supplying the pLenti*-Penk1-Cre* and pLenti*-Penk1-GFP* vectors used in this study. We would also like to EPFL BiOP imaging facility for technical assistance. This work was supported by grants from the Swiss National Science Foundation (SNSF) (176206; NCCR Synapsy: 51NF40-158776 and −185897) to C.S.. S.G. was supported by Eurotech EU Horizon 2020 (No. 754462) and independent postdoctoral funding from the AXA Research Funds. E.R-F. was supported by European Union’s Horizon 2020 research and
innovation programme under the Marie Sklodowska-Curie grant agreement N° 895562.

## Authors Contributions

Concept development and experimental design, S.G., S.A., and C.S; Animal surgeries, E.G., S.G., A.C., J.G.; Behavioral tests and analyses, E.G., S.G.; *Ex vivo* analyses, E.G., S.G., E.R-F.; Electrophysiological recordings and analyses, S.A.; Provision of key resources, B.S., A.Z.; Writing, S.G., S.A., C.S.. All authors discussed the results and edited and approved the manuscript.

## STAR METHODS

### LEAD CONTACT AND MATERIALS AVAILABILITY

Further information and requests for resources and reagents should be directed to and will be fulfilled by the Lead Contact, Carmen Sandi.

#### Materials availability

This study did not generate new unique reagents.

### EXPERIMENTAL MODEL AND SUBJECT DETAILS

Animal care and experimental procedures were conducted in accordance with the Swiss Federal Guidelines for Animal Experimentation and were approved by the Cantonal Veterinary Office Committee for Animal Experimentation (Vaud, Switzerland).

#### Rats

Adult male Wistar rats (Charles River, L’Arbresle, France) were individually housed in polypropylene cages (57 × 35 × 20 cm) with abundant pine bedding in a temperature-(23°C) and light-(0700-1900 h) controlled room. All animals had *ad libitum* access to standard food and water. Upon arrival to the facility, animals were allowed to habituate to the vivarium for one week and were then handled for 2 min/d during 3 days prior to the start of all experiments. All behavioral manipulations were performed during the light phase.

#### Mice

All mice used for these studies were male and were on the C57BL/6 background. The generation Mfn2^loxp/loxp^ (Mfn2^f/f^) (Chen et al., 2007) and Adora2a-Cre (Gong et al., 2007) mice have been previously reported. To generate Adora2a-specific constitutive Mfn2 knock-out mice (hereafter, Mfn2^Adora2a^), Adora2a-Cre were crossed with Mfn2^f/f^ mice. Mice were maintained on a 12:12h light– dark cycle with free access to water and standard chow, unless noted otherwise.

### Genotyping

All *Mfn2*^*f/f*^ and *Mfn2*^*Adora2a*^ mice were genotyped before weaning (from ear punches) by polymerase chain reaction (PCR) using the following primers: Mfn2 FW: 5’ TTT GGA AGT AGG CAG TCT CCA 3’; Mfn2 RV: 5’ CAG GCA GCA CTG AAA AGA GA 3’; Adora2a-Ctrl FW: 5’ GTT ACC TAT TGA ACG CCC TAC 3’; Adora2a-Ctrl RV: 5’ GCC AAC AAA GTT TAG ATG TAT CTA AGG 3’; Adora2a-Cre FW: 5’ CCG GTG AAC GTG CAA AAC AGG CTC 3’; Adora2a-Cre RV: 5’ GGC AGA TGG CGC GGC AAC AC 3’.

### Constructs and virus

Plasmid encoding Pdyn-Cre (Darvas and Palmiter, 2015) was a gift from Dr. Martin Darvas, University of Washington, Seattle, USA. The construct was packaged in AAV2 serotype at the EPFL viral core. control AAV, we used the AAV2-SYN-eGFP-WPRE vector. AAV2 vector titers were 2.2E+11 VG/ml as determined by PCR. The plasmids encoding pLenti-Penk1-Cre and pLenti-Penk1-GFP (Zhang et al., 2015) were a gift from Dr. Fan Wang, Duke University, North Carolina, USA. The titers were 8’811µg/ml and 5’464µg/ml as determined by p24 antigen. The pAAV9-hSyn1-Myc-hMFN2 vector was a gift from Dr. Bernard Schneider, EPFL, and the titer was 10^9^ VG/µl.

### Surgery

#### Rats

For Mfn2 overexpression, pAAV9-hSyn1-Myc-hMFN2 virus was delivered bilaterally into the NAc with two injections (1.3 and 2.5⍰mm posterior to bregma, 1.0 and 1.5⍰mm from midline, 7.0 ventral from skull) at a volume of 0.8⍰μl with a constant flow rate of 0.1⍰μl/min. The injectors were left in site for 5 min after the end of the actual injection. After removing the injectors, animals were treated with paracetamol (500⍰mg/700⍰ml Dafalgan, Bristol-Myerts Squibb, Agen, France) via the drinking water for seven days after the surgery. Behavioral experiments commenced 4 to 5 weeks following surgery.

#### Mice

Mice were anesthetized by inhalation of a 3% isoflurane mix in O_2_ gas (produced by the CombiVet animal gas anesthesia system; Rothacher Medical, Switzerland) and maintained under inhalation of 1 to 1.3% isoflurane in O_2_. Analgesia (buprenorphine at 0.1 mg/kg) was injected subcutaneously. Mice were then placed in a stereotactic apparatus (Kopf Instruments, model 940, USA) on a heating pad (Harvard Instruments, USA). Following exposure of the skull, a hole was drilled on each side with a 0.5-mm bore (Komet Dental, Germany) using a Tech2000 drill handpiece (Ram Products Inc, USA). For Mfn2 knockdown, either AAV2-Pdyn-Cre or pLenti-Penk1-Cre or corresponding control viral vectors were delivered into the NAc (distance from bregma, AP: 1.6 mm; ML: ±0.8 mm; DV: − 4.7 mm). The injectors were left in site for 10⍰min after the end of the actual injection. After removing the injectors, animals were treated with paracetamol (500⍰mg/700⍰ml Dafalgan, Bristol-Myerts Squibb, Agen, France) via the drinking water for 5 days after the surgery. Behavioral experiments commenced 4 weeks following surgery.

### Behavioral testing

#### Elevated plus-maze (EPM) test

the maze consisted of a central platform elevated from the ground from which two opposing open and two opposing closed arms emanated, as described previously (Gebara et al., 2021). Lighting was maintained at 15-16 lx on the open arms and 5-7 lx in the closed arms. The experimental animal was gently placed in the center of the maze looking towards a closed arm and allowed to move undisturbed for 5 min. After each trial, the arms were cleaned with 7% ethanol and dried. Video tracking of the animal’s location was performed by a camera fixed above the arena. Ethovision tracking system (Ethovision 11.0 XT, Noldus, Information Technology) was used to calculate the percentage of time spent in the open arms, which was taken as an indicator of anxiety. Rats were classified as high-(HA, ≤5% open arm duration), intermediate-(IA, 5-20% open arm duration) or low-anxious (LA, ≥20% open arm duration).

#### Open field (OF) and novel object (NO) tests

The experimental animal was placed in the corner of a rectangular arena (for rats, as described by (Gebara et al., 2021) and for mice as described by (Morató et al., 2022) and left to freely explore for 10 min (OF test phase). Afterwards, a new object was placed in the centre of the arena for 5 min (NO test phase). Lighting was maintained at 8-10 lux on the center of the arena. After each trial, the arena was cleaned with 7% ethanol and dried. Video tracking of the animal’s location was performed by a camera fixed above the arena. The percentage of time spent in the center of the open field was taken as an indicator of anxiety (OF test phase) or reactivity upon novelty (NO test phase).

#### Social preference test

Social exploration in rats was evaluated as previously described (van der Kooij et al., 2018). Rats are introduced in a three-chambered box (35 × 30 cm for side chambers, 35 × 20 cm for middle chamber), which they are allowed to explore for 10 minutes. Within the testing time, the experimental rats can choose between an interaction with a stranger rat (juvenile, Wistar) or an object that are enclosed in a wire cylinder in opposite side chambers. The time spent sniffing each cylinder was manually scored by an experimenter blind to the treatments to evaluate the level of preference for the unfamiliar juvenile compared with the object.

#### Social dominance test (in rats)

Pairs were matched for weight or different anxiety profile. For each dyad, the rats were considered equal in their probability (= 50%) to become dominant or subordinate during their encounter. The social competition was performed as previously described (Hollis et al., 2015; van der Kooij et al., 2018), including *a priori* exclusion of any pairs exhibiting 10 s or less of offensive behavior from analysis.

#### Social dominance test (in mice)

Social dominance in mice was evaluated as described previously (Wang et al., 2011) with some modifications. Mfn2^f/f^ mice (injected with GFP or Cre targeting pDyn/D1- or Enk/D2-MSNs) were matched for weight and were pair-housed for two weeks prior to the dominance tube test. The tube test apparatus consisted of a transparent Plexiglas tube with 30 cm length and 3 cm inner diameter. This diameter is a size just sufficient to allow adult mice to pass through without reversing the direction. The test consisted of two phases: a training phase (2 days) and a test phase (3 days). During the training phase, each mouse was trained to pass through the tube. Mice were not permitted to escape backward out of the tube. During the test phase, two mice were released simultaneously into the opposite ends and care was taken to ensure that they met in the middle of the tube. The mouse that first retreated from the tube ending with all four paws on the outside within 2 min was designated the loser of the trial. Each pair was tested in three encounters during 3 consecutive days. Each mouse was ranked by its winning scores (between 0 and 3, with 3 indicating a win). Trials were scored by an individual blind to the genotypes.

#### Warm spot test

The warm spot test was adapted from a protocol previously described (Zhou et al., 2017). Briefly, a rectangular plastic cage (28 × 20 cm) was cooled on ice until the floor reached 0-4°C. Under one corner of the cage, a waterproof heating pad was placed to locally heat the floor to 32°C. The nest was 5 cm in diameter and just big enough to permit the stay of only one adult mouse. Temperatures were measured by an infrared thermometer. Twenty min prior to trial initiation, mice were transferred from their home cage to an empty cage maintained on ice to cool down, and then transferred to the test cage where they competed for the nest on the warm spot. For each trial, behavior was tracked for 10 min and occupation of warm nest was measured.

#### Wire hang test

To evaluate motor function and muscle strength, each mouse was hanged from an elevated wire cage top. The mouse was placed on the cage top, which was then inverted and suspended above the home cage; the latency to when the animal falls was recorded. This test was repeated for three times.

### Body composition

Whole body composition was determined by nuclear magnetic resonance (NMR)-based technology (EchoMRI, Echo Medical Systems, Houston, TX, USA). Each mouse was placed briefly (approximately 1 min, no anesthesia required) in the EchoMRI machine where lean and fat mass were measured. Whole body fat and lean content are expressed as a percentage of total body weight.

### Brain region microdissection

Mice were decapitated, and brain was extracted and snap-frozen in isopentane at −45⍰°C and stored at −80⍰°C until further processing. The brains were sectioned on a freezing (−20 °C) cryostat (Leica) and 150⍰μm-thick slices were mounted onto SuperFrost™ Plus Microscope Slides (Thermo Fisher Scientific, USA). NAc from GFP and Cre-injected mice was dissected using 0.5-1.5 mm tissue punches (Harris UniCore, USA) according to the atlas coordinates (Paxinos and Franklin, 2001). Tissue was collected in RNAase-free tubes and maintained in a −80⍰°C freezer until further processing.

### RNA extraction, cDNA synthesis, and qPCR

For RNA extraction, animals were decapitated with a guillotine and brains were flash frozen in ice-cold isopentane and then stored at − 80 °C. The NAc was dissected for gene expression analyses by tissue punching 200 μm slices sectioned on a freezing cryostat. We used a core punch sampler (Harris, Uni-core, 1.5 mm for mice and 2.0 mm for rats) to harvest bilateral NAc tissue on both sides. Tissue samples were placed immediately on dry ice and stored at − 80°C until RNA or protein extraction. Total RNA from the tissue samples was purified using a RNAqueous Total RNA Isolation kit (Ambion, AM1912) and quantified by NanoQuant Plate and Spark reader (Tecan). 200 ng of total RNA was then reverse transcribed to cDNA with qScript (Quanta Biosciences, 95048-500). The resulting cDNA was diluted to 2 ng/μl. For each qPCR, 1.5 μl of cDNA was combined with 5 μl of Power SYBR Green PCR Master Mix (Thermo Fisher Scientific, 4368708), forward/reverse primers (3.5 μl total), and 1 μl of water. The qPCR reactions were performed in triplicates in an ABI Prism 7900 Sequence Detection System (Applied Biosystems). The standard cycling conditions were 95°C for 10 min, followed by 40 cycles of 95°C for 15 s, and 60°C for 1 min. Melt curves were generated at the end of the regular qPCR cycles. Analysis was performed using the ΔΔC(t) method. Samples were normalized to the EEF1 level. The gene specific PCR primer sequences are shown in the key resources table.

### Golgi staining and Sholl analysis

Animals were rapidly decapitated, and brains were removed quickly from the skull. After rinsing with PBS, the brains were stained with the FD Rapid GolgiStain™ kit (FD NeuroTechnologies, Ellicott City, MD, USA). They were first immersed in the impregnation solution (A and B) which was replaced after 6–12 h, and were then kept in dark for 15–16 days. Afterwards, the brains were put in Solution C, which was replaced after 24 h and kept in dark for the next 48–60 h. Cryomicrotome (Microm Thermo Scientific, Walldorf, Germany) was used to cut 200 μm thick slices. Slices were mounted on a gelatin-coated microscope slide, stained, dehydrated and coverslipped with Permount. Tissue preparation and staining were all done by the same person following the FD Rapid GolgiStain™ kit manufacturer’s protocol. Medium spiny neurons were reconstructed using a Leica DM500 equipped with x-y motorized stage with a 63x objective (Ludl Electronic Products Ltd, Hawthorne, NY, USA). Sholl analysis was performed to measure dendritic length and number of intersections per concentric circles starting from the point at the centroid of the cell body.

### Hematoxylin-eosin (HE) staining

HE staining was performed as described previously (Hollis et al., 2015). Briefly, brain slices (35 μm) were mounted on a slide and dehydrated using increasing steps of EtOH. Sections were incubated with hematoxylin for 5 min, gently rinsed in tap water for 10 min, and stained in eosin for 2 min. Sections were rinsed in distilled water, dehydrated, and mounted. Images were captured using a brightfield slide scanner (Olympus Slide Scanner VS120-L100).

### Proximity ligation assay

Fixation and blocking were performed as described for immunocytochemistry. To label VDAC-IPR3 interacting dots, slices were incubated overnight at 4°C with rabbit anti-IP3R and mouse anti-VDAC1 primary antibodies in the antibody reagent buffer provided in the proximity ligation assay (PLA) kit (Duolink, Sigma-Aldrich). Slices were then washed, and incubated with the anti-mouse minus and anti-rabbit plus probes in the antibody reagent buffer for 1 h at 37°C. If the antigens of interests are closer than 40 nm, connector oligonucleotides can hybridize with the probes and, after ligation, the signal is enhanced by rolling circle amplification. Coverslips were mounted in Vectashield mounting medium containing DAPI. We used the red Duolink detection fluorophore. The fluorescent spots corresponding to the VDAC1-IP3R1 interactions were imaged using a confocal microscope under a 40x objective with 2x digital zoom and analyzed using Fiji software using analyze particles plugins. The number of particles or dots was normalized by the number of nuclei.

### Mitochondrial respiration

Mice were sacrificed by rapid decapitation and the NAc was rapidly dissected out, weighed, and placed in a Petri dish on ice with 2 ml of relaxing solution (2.8 mM Ca_2_K2EGTA, 7.2 mM K_2_EGTA, 5.8 mM ATP, 6.6 mM MgCl_2_, 20 mM taurine, 15 mM sodium phosphocreatine, 20 mM imidazole, 0.5 mM DTT and 50 mM MES, pH = 7.1) until further processing. Tissue samples were then gently homogenized in ice-cold respirometry medium (MiR05: 0.5 mM EGTA, 3mM MgCl_2_, 60 mM potassium lactobionate, 20 mM taurine, 10 mM KH_2_PO_4_, 20 mM HEPES, 110 mM sucrose and 0.1% (w/v) BSA, pH=7.1) with an eppendorf pestle. Then, 2 mg of tissue were used to measure mitochondrial respiration rates at 37°C using high resolution respirometry (Oroboros Oxygraph 2K, Oroboros Instruments, Innsbruck, Austria), as previously described (Hollis et al., 2015). A substrate-uncoupler-inhibitor titration (SUIT) protocol was used to sequentially explore the various components of mitochondrial respiratory capacity. A non-phosphorylating resting state (Leak) was obtained after adding a mixture of malate (2 mM), pyruvate (10 mM) and glutamate (20 mM). In order to measure the respiration due to oxidative phosphorylation (OXPHOS), we added substrates for the activation of specific complexes. Thus, oxygen flux due to complex I activity (Complex I) was quantified by the addition of ADP (5 mM), followed by the addition of succinate (10 mM) to subsequently stimulate complex II (CI & CII). We then uncoupled respiration to examine the maximal capacity of the electron transport system (ETS) using the protonophore, carbonylcyanide 4 (trifluoromethoxy) phenylhydrazone (FCCP; successive titrations of 0.2 µM until maximal respiration rates were reached). We then examined consumption in the uncoupled state due solely to the activity of complex II (CII) by inhibiting complex I with the addition of rotenone (0.1 µM). Finally, electron transport through complex III was inhibited by adding antimycin (2 µM) to obtain the level of residual O_2_ consumption (ROX) due to oxidative side reactions outside of mitochondrial respiration. The O_2_ flux obtained in each step of the protocol was normalized by the wet weight of the tissue sample used for the analysis and corrected for ROX.

### ATP quantification

ATP was measured with the CellTiter-Glo Luminescent Cell Viability Assay (Promega), with a few minor modifications. Freshly dissected NAc tissue was placed in 2 ml of relaxing solution on ice. Tissue samples were then diluted 10x in a tricine buffer solution (40 mM Tricine, 3 mM EDTA, 85 mM NaCl, 3.6 mM KCl, 100 mM NaF and 0.1% saponin, pH 7.4; Sigma-Aldrich). ATP content was determined enzymatically with luciferase in a white 96-well plate. In the presence of ATP, Mg^2+^, and oxygen, luciferin is oxygenated by luciferase into oxyluciferin. This reaction emits light which is proportional to the amount of ATP in the sample. A converting solution (100 mM Tricine, 100 mM MgSO_4_, 25 mM KCl) was added to tissue samples and allowed to incubate at room temperature for 5 min. After incubation, a MgCl_2_ solution (4 mM tricine and 100 mM MgCl_2_) was added to the samples, followed by, 35 μl of CellTiter-Glo reagent (G7571, Promega). Additionally, each 96-well plate contained a series of 10-fold dilutions (1 µM–10 nM) of an ATP standard (Sigma), in order to generate a standard curve for each assay. Luminescence was immediately detected with a luminometer (Safire 2, Tecan). Luminescence was measured kinetically via 40 cycles taken at 1 min intervals. At least 4 points at the steady-state were taken to generate an average maximum luminescence for each sample. ATP was calculated using the standard curve to determine the concentration of ATP.

### Immunofluorescence

Coronal mouse brain sections were rinsed three times for 10⍰min in 0.1 M PBS followed by a blocking step of 11h incubation in blocking solution (0.1 M PBS containing 3% bovine serum albumin (Sigma Aldrich, Buchs, Switzerland) and 0.3% Trition-X100). Primary antibodies against GFP (abcam, 1:1000 dilution), NeuN (EMD Millipore, 1:1000 dilution) or cleaved caspase 3 (Cell Signaling, 1:200 dilution) were diluted in blocking solution, and sections were incubated overnight at 4°C with gentle shaking. Sections were then washed three times in PBS and incubated with secondary antibodies (goat anti-chicken Alexa Fluor 488, abcam, 1:500 dilution, or donkey anti-rabbit Alexa 568, Thermo Fisher Scientific, 1:500 dilution) for 2⍰h at room temperature. After three rinses in PBS, sections were incubated for 2⍰min with 4,6-diamidino-2-phenylindole (DAPI; Sigma, D9542), washed again three times in PBS and then mounted in Vectashield (Vector Labs). GFP, NeuN or caspase 3 expression was assessed in the NAc using a LSM 710 laser-scanning confocal microscope (Carl Zeiss) imaged using a 10x, 20x, or 40x oil immersion objective with a 1.0 digital zoom. Images were quantified for number of NeuN positive cells or fluorescence intensity for caspase-3 activation using ImageJ (NIH, USA). At least two sections from each animal were measured and averaged to generate one value per hemisphere per animal for each group.

### Hormone analyses

Trunk blood was collected and centrifuged at 9400g at 4°C for 4 min to obtain plasma. Plasma samples were prepared according to manufacturer’s instructions to measure corticosterone and testosterone concentrations using an ELISA kit (Enzo Life Sciences, ADI-901-097 for corticosterone, ASI-901-065 for testosterone). Levels were calculated using a standard curve method.

### *Ex vivo* electrophysiology

Enk^GFP^ and Enk^Cre^ mice were anesthetized with isoflurane and decapitated. The brain was quickly removed and placed in oxygenated (95% O_2_ / 5% CO_2_) ice-cold modified artificial CSF (ACSF), containing (in mM): 105 sucrose, 65 NaCl, 25 NaHCO_3_, 2.5 KCl, 1.25 NaH_2_PO_4_, 7 MgCl_2_, 0.5 CaCl_2_, 25 glucose, 1.7 L(+)-ascorbic acid. Coronal slices (250 µm thick) containing the ventral striatum were cut using a vibrating tissue slicer (Campden Instruments) and incubated in standard ACSF containing (in mM): 130 NaCl, 25 NaHCO_3_, 2.5 KCl, 1.25 NaH_2_PO_4_, 1.2 MgCl_2_, 2 CaCl_2_, 18 glucose, 1.7 L(+)-ascorbic acid, and complemented with 2 Na-pyruvate and 3 myo-inositol. The incubation temperature was

∼35°C during the first hour. In the recording chamber, slices were superfused with oxygenated standard ACSF at room temperature (for synaptic current recordings), or 30-32°C for cell firing recordings. GFP-positive neurons in the NAc shell were patched in the whole-cell configuration with borosilicate pipettes (3-4 MΩ) filled with (in mM): 120 CsGluconate, 10 CsCl, 10 HEPES, 10 phosphocreatine, 5 EGTA, 4 Mg-ATP for synaptic current recordings, or 135 mM KGluconate, 10 mM KCl, 10 mM HEPES, 0.2 mM EGTA, 1.5 mM Mg-ATP, 0.2 mM Na-GTP for cell firing recordings (290-300 mOsm, pH 7.2-7.3). Miniature excitatory postsynaptic currents (mEPSCs) were recorded at the holding potential of −60 mV in the presence of the GABA_A_R blocker picrotoxin (0.1 mM) and the Na^+^ channel blocker Tetrodotoxin (0.001 mM). Synaptic currents were acquired for 5 min starting from >8 min after the establishment of the whole-cell configuration, to ensure proper diffusion of the intracellular solution. To elicit neuronal firing, cells were held at −60 mV in current clamp configuration with direct current injections, and depolarization was provided by 5-s long current ramps of increasing magnitude (50 pA-steps of maximal current). Electrophysiological data were acquired through a Digidata1550A digitizer. Signals were amplified through a Multiclamp700B amplifier (Molecular Devices), sampled at 20 kHz and filtered at 10 kHz using Clampex10 (Molecular Devices). Data were analysed using Clampfit10 (Molecular Devices). For detection of mEPSCs, traces were filtered at 1 kHz and analysed with Easy Electrophysiology v2.3 (Easy Electrophysiology Ltd., UK) using the template detection method and an amplitude threshold of 5 pA. Detected events were verified by visual inspection. To construct cumulative frequency plots, the first 200 events recorded in each cell were considered.

### Statistics

Statistical analyses were performed in Prism v.7.0 or v.8.0 (GraphPad Software). For experiments that included two groups, the results were analyzed using two-tailed Student’s t-tests or two-tailed Mann-Whitney U test (in cases where the data did not meet assumptions for parametric statistics). For cumulative frequency plots, Kolmogorov-Smirnov test was used between groups to calculate differences between distributions. P-values < 0.05 were considered to be significant and p-values < 0.1 a trend. # indicates p < 0.1, * indicates p < 0.05, ** indicates p < 0.01, and *** indicates p < 0.001. In graphs, individual points represent single subjects in all behavioral experiments, cells in morphology experiments, sections in anatomical analyses. For the Sholl analysis, parametric tests were used: two-way ANOVA with repeated-measures was performed, followed by a post hoc analysis. All data are presented as mean ± SEM in bar graphs and min-to-max box-and-whisker plots. The statistical details of each experiment can be found in Table S1.

