## Supplementary material for "Mitofusin-2 in nucleus accumbens D2-MSNs regulates social dominance and neuronal function": Suppl Info

**Supplemental Information**

**Figures S1 to S4**

**Table S1**

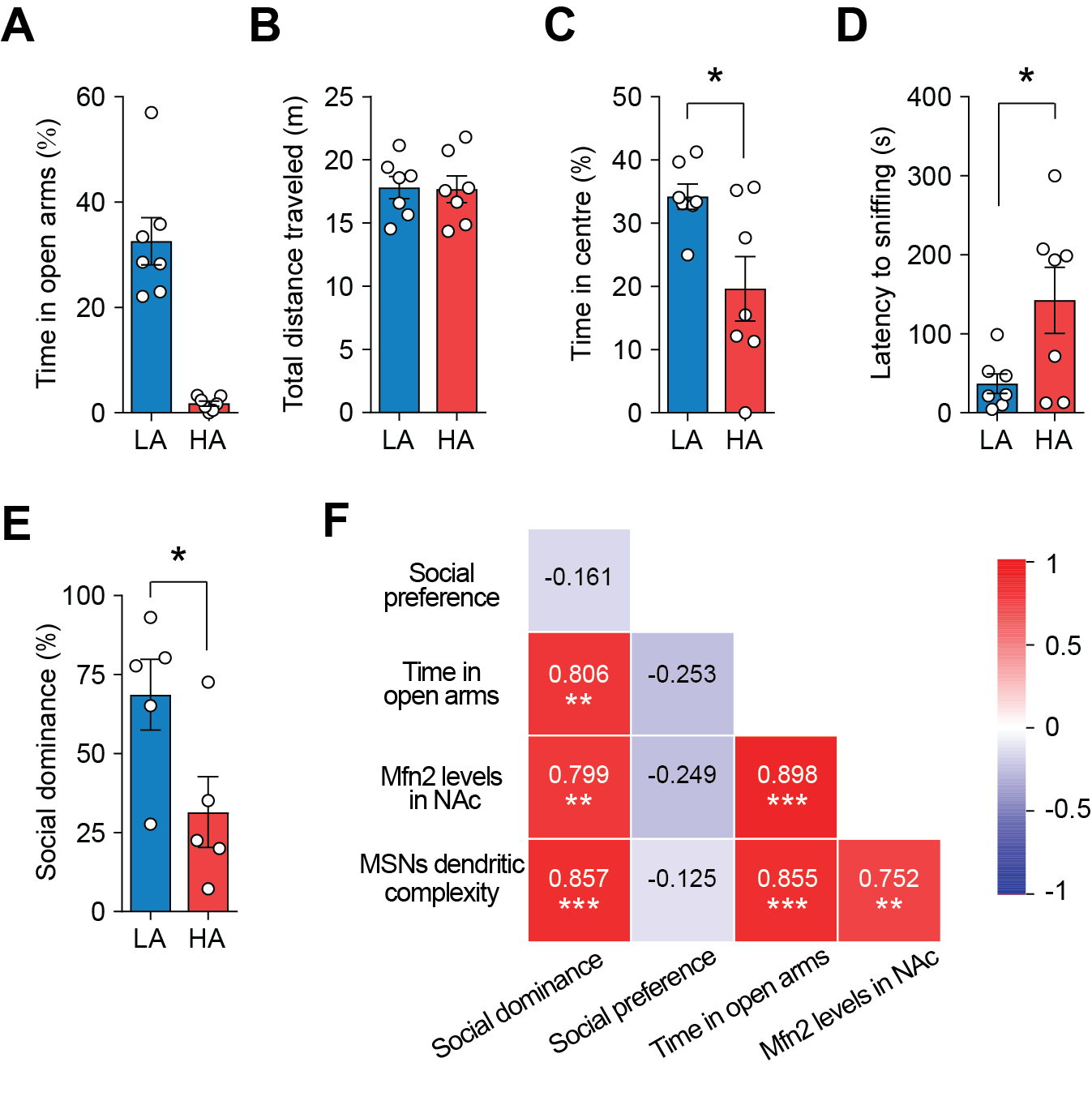

**Figure S1. Additional behavioral and cellular analyses in LA and HA rats. Related to Figure 1.**

(A) Time spent in the open arms of the elevated plus maze (EPM) of rats classified as low anxious (LA) or highly anxious (HA). (B) Total distance moved by LA and HA rats in the EPM. (C) Time spent by LA and HA rats in the center of the open field (OF). (D) Latency to explore the novel object (NO) placed in the center of the OF. (E) HA rats showed decreased social dominance when confronted with LA rats. (F) Correlation plot showing positive relationship between social dominance, time in the open arms of the EPM, *Mfn2* gene expression in the NAc and dendritic complexity of NAc MSNs. Social preference did not correlate with any of the previous parameters (Pearson correlation coefficients are reported in the color-coded cells). Data are mean ± SEM in bar graphs. Circles in the bar graphs represent single observations. * p < 0.05, ** p < 0.01, *** p < 0.001. Exact statistics can be found in Table S1.

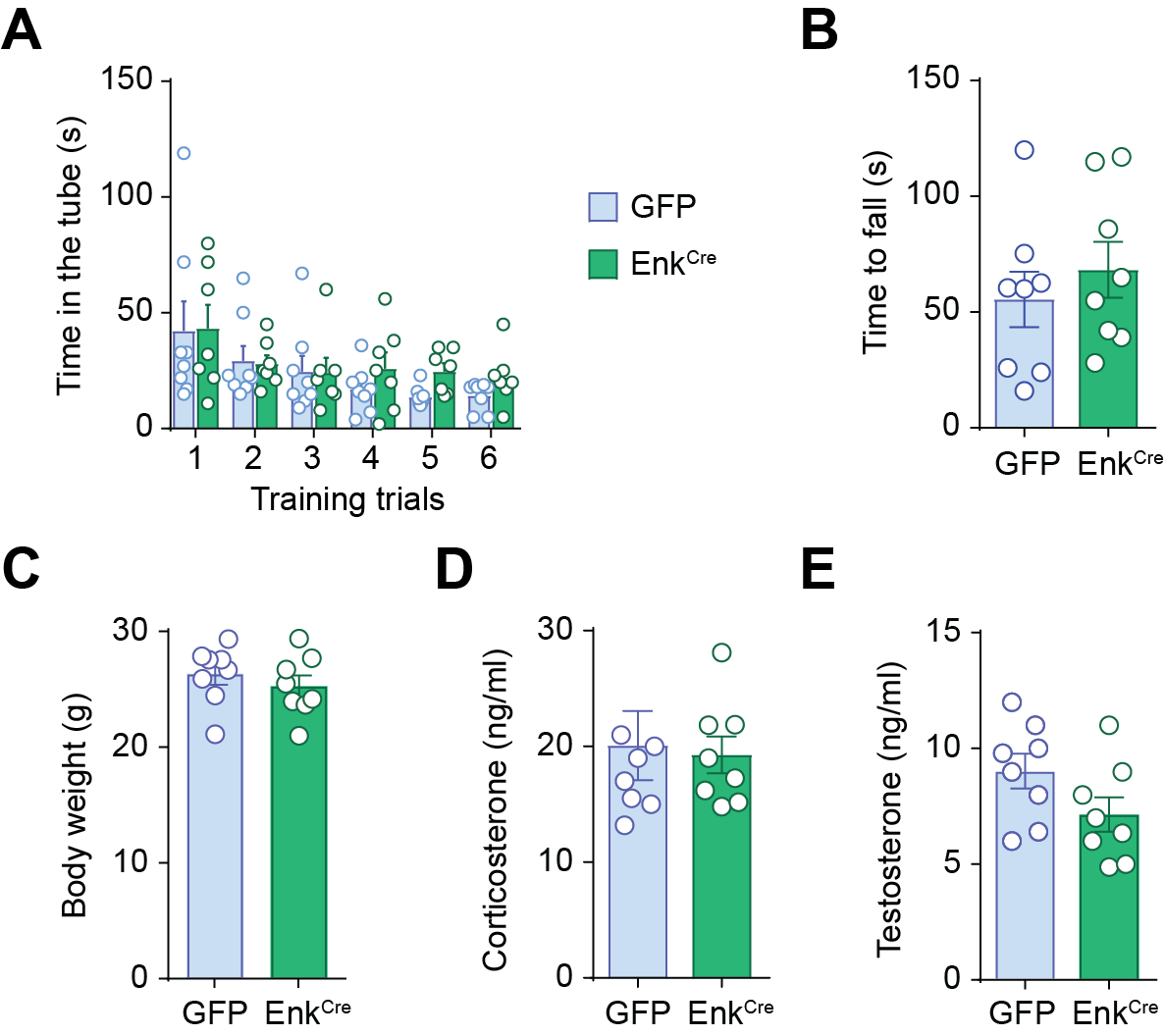

**Figure S2. Additional analyses of behavioral parameters and somatic endpoints in GFP and Enk^Cre^ mice. Related to Figure 2.**

(A) Enk^Cre^ and GFP mice readily learnt to come out of the tube during the training trials. (B) Muscle strength was not impaired in the Enk^Cre^ mice, as indicated by the wire hang test. (C) Body weight was not affected in the Enk^Cre^ mice. (D-E) GFP and Enk^Cre^ mice showed similar plasma corticosterone and testosterone levels. Data are mean ± SEM. Circles in the bar graphs represent single observations. Exact statistics can be found in Table S1.

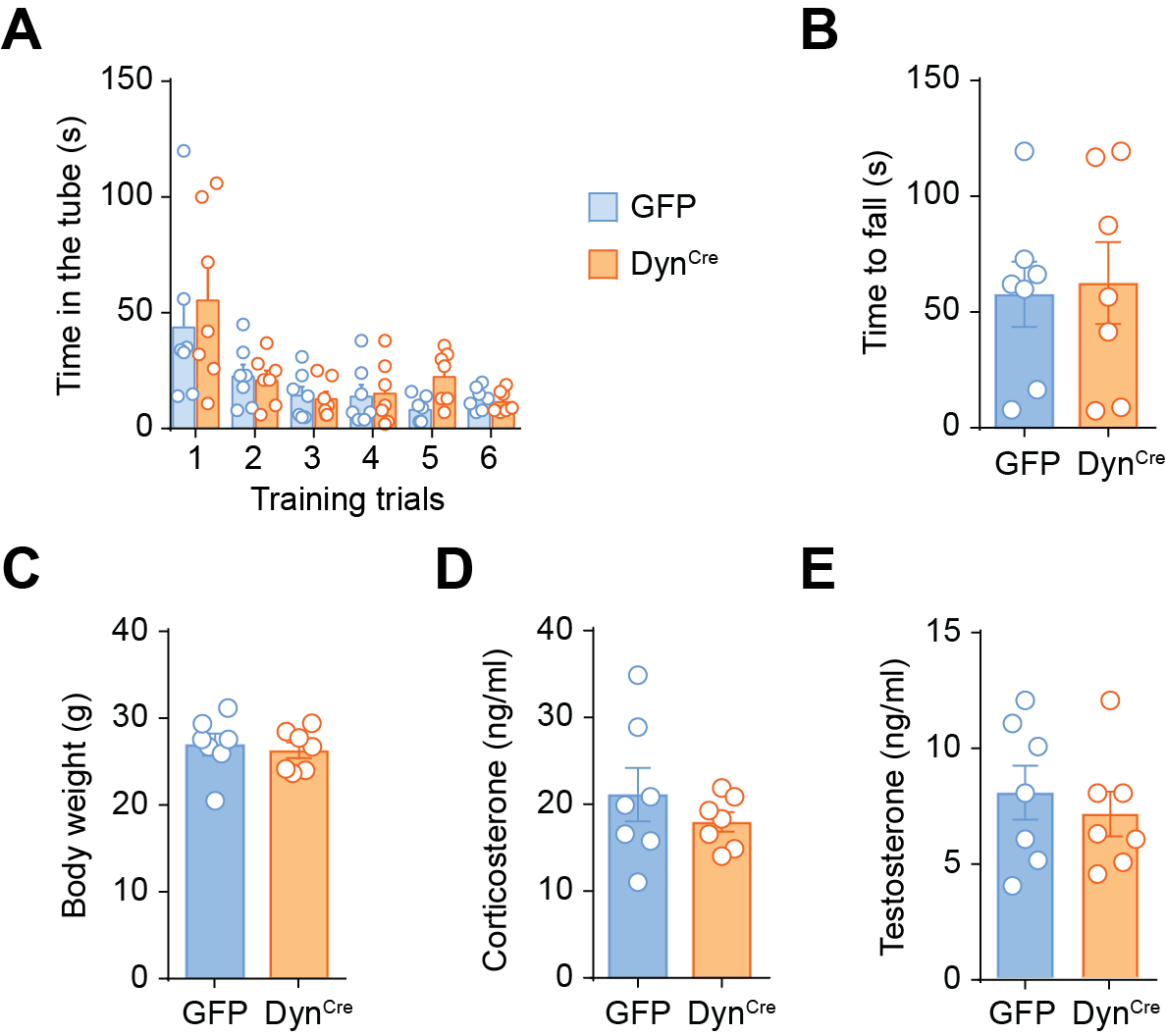

**Figure S3. Additional behavioral analyses and somatic endpoints in GFP and Dyn^Cre^ mice. Related to Figure 3.**

(A) GFP and Dyn^Cre^ mice readily learnt to come out of the tube during the training trials. (B) Muscle strength was not impaired in the Dyn^Cre^ mice, as indicated by the wire hang test. (C) Body weight was not affected in the Dyn^Cre^ mice. (D-E) GFP and Enk^Cre^ mice showed similar plasma corticosterone and testosterone levels. Data are mean ± SEM. Circles in the bar graphs represent single observations. Exact statistics can be found in Table S1.

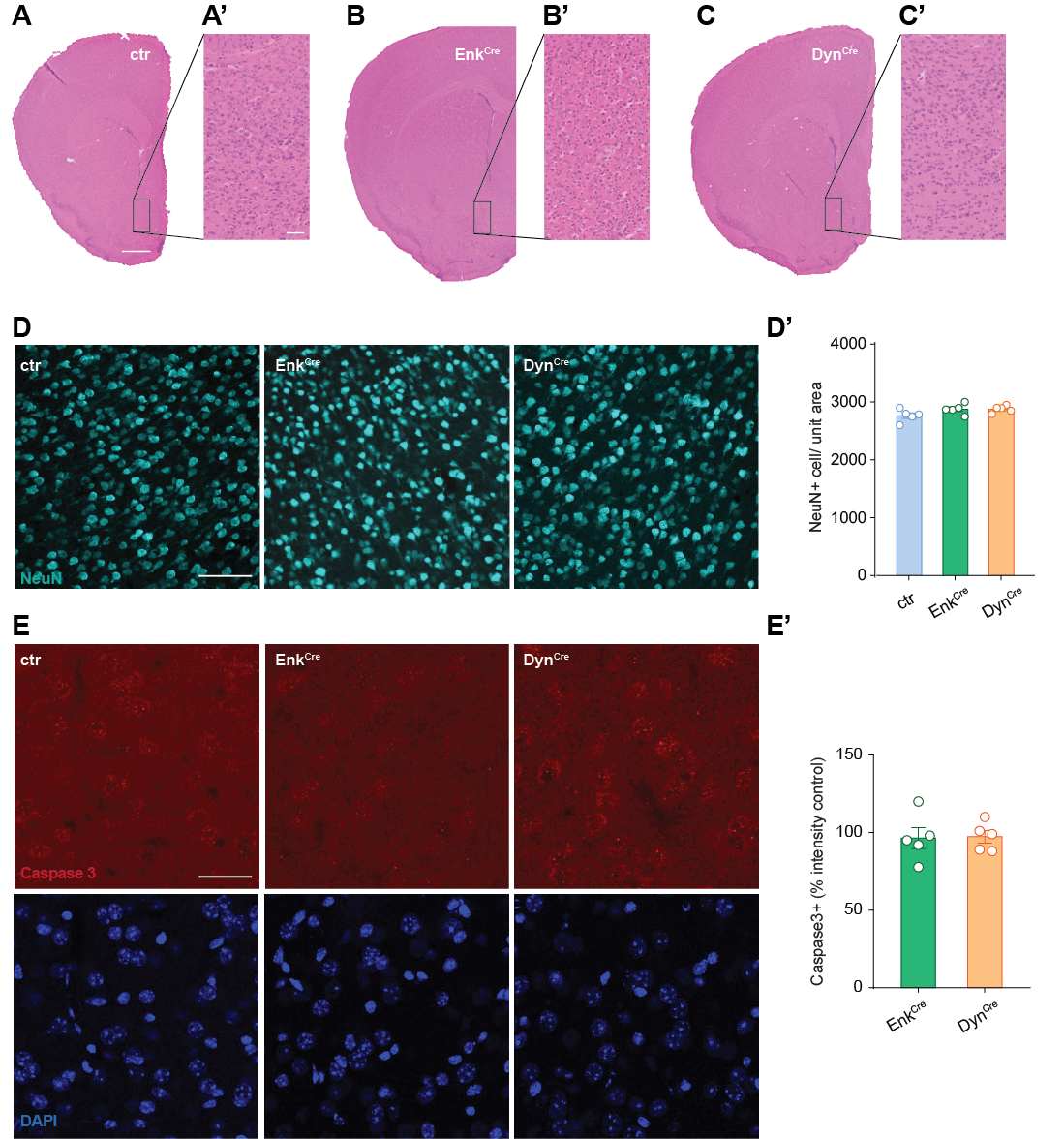

**Figure S4. Mfn2 knockdown in Dyn or Enk neurons does not affect neuronal viability in the NAc**. **Related to Figures 4-5.**

(A-C’) Representative low (A, B, C) and high (A’, B’, C’) magnification images of Hematoxylin-eosin (H&E) staining of control (ctr) or Cre-injected NAc regions revealed intact patterns of NAc morphology in Enk^Cre^ or Dyn^Cre^ mice. Scale bar = 1 mm in A applies to B and C, and scale bar = 50 µm in A’ applies to B’ and C’. (D-D’) Representative NeuN immunolabeled sections after microinjection of control (ctr) or Enk^Cre^ or Dyn^Cre^ viruses. Scale bar = 100 µm. Quantification of NeuN positive cells shows that NeuN immunoreactivity remained intact in the NAc region following microinjection of the respective Cre viruses and Mfn2 knockdown, indicating that neuronal viability was not affected. (E-E’) Representative cleaved caspase 3 (red) immunolabeled sections after intra-NAc injection of control (ctr), Enk^Cre^ or Dyn^Cre^ viruses. Scale bar = 50 µm. There were no effects of Mfn2 knockdown on apoptotic cell death, as evidenced by the amount of cleaved-caspase 3. Data are mean ± SEM. Circles in the bar graphs represent single observations. Exact statistics can be found in Table S1.

| **Figure** | **Measure** | **Subjects** | **Analysis** | **Factors** | **Statistics Value** | **p Value** | **Post hoc test** | **Data normally distributed** |
| --- | --- | --- | --- | --- | --- | --- | --- | --- |
| 1C left | Social dominance (%) | LA (n=5 rats) vs Mfn2OE (n=5 rats) | unpaired t test, two tailed |  | t(8)=3.493 | p=0.0082 |  | YES |
| 1C right | Social dominance (%) | HA (n=5 rats) vs Mfn2-OE (n=5 rats) | unpaired t test, two tailed |  | t(7.31)=2.799 | p=0.0254 |  | YES |
| 1D | Social preference (%) | LA (n=6 rats) vs HA (n=6 rats) vs Mfn-2OE (n=6 rats) | One-way ANOVA |  | F(2,15)=0.5001 | p=0.6162 |  | YES |
| 2B | Relative expression of Mfn2 mRNA | Enk^Cre^ (n=8 mice) vs GFP (n=8 mice) | unpaired t test, two tailed |  | t(14)=10.96 | p<0.0001 |  | YES |
| 2D | # Wins across days | Enk^Cre^ (n=8 mice) vs GFP (n=8 mice) | unpaired t test, two tailed |  | t(14)=5.65 | p<0.0001 |  | YES |
| 2E | # Wins per day | Enk^Cre^ (n=8 mice) vs GFP (n=8 mice) | Two-way RM ANOVA per mouse | day x  genotype | interaction F(2, 28)=0.58  day F(2, 24)=0.02 genotype F(1,14)=34.18 | p=0.5659  p=0.9727  p<0.0001 | Bonferroni's multiple comparisons test | YES |
| 2F | # Pushes | Enk^Cre^ (n=8 mice) vs GFP (n=8 mice) | unpaired t test, two tailed |  | t(14)=5.49 | p<0.0001 |  | YES |
| 2G | Push duration | Enk^Cre^ (n=8 mice) vs GFP (n=8 mice) | unpaired t test, two tailed |  | t(14)=4.46 | p=0.0005 |  | YES |
| 2H | # Retreats | Enk^Cre^ (n=8 mice) vs GFP (n=8 mice) | unpaired t test, two tailed |  | t(14)=4.93 | p<0.0001 |  | YES |
| 2I | Time retreating (%) | Enk^Cre^ (n=8 mice) vs GFP (n=8 mice) | unpaired t test, two tailed |  | t(14)=6.50 | p<0.0001 |  | YES |
| 2K | Time in the warm spot (%) | Enk^Cre^ (n=8 mice) vs GFP (n=8 mice) | unpaired t test, two tailed |  | t(14)=3.14 | p=0.007 |  | YES |
| 2L | Time in the center of OF (%) | Enk^Cre^ (n=8 mice) vs GFP (n=8 mice) | unpaired t test, two tailed |  | t(14)=0.38 | p=0.70 |  | YES |
| 2M | Distance moved in the OF (m) | Enk^Cre^ (n=8 mice) vs GFP (n=8 mice) | unpaired t test, two tailed |  | t(14)=0.64 | p=0.52 |  | YES |
| 2N | Time sniffing NO (%) | Enk^Cre^ (n=8 mice) vs GFP (n=8 mice) | unpaired t test, two tailed |  | t(14)=1.36 | p=0.19 |  | YES |
| 3B | Relative expression of Mfn2 mRNA | Dyn^Cre^ (n=7 mice) vs GFP (n=7mice) | unpaired t test, two tailed |  | t(12)=7.73 | p<0.0001 |  | YES |
| 3C | # Wins across days | Dyn^Cre^ (n=7 mice) vs GFP (n=7mice) | unpaired t test, two tailed |  | t(12)=0.64 | p=0.55 |  | YES |
| 3D | # Wins per day | Dyn^Cre^ (n=7 mice) vs GFP (n=7mice) | Two-way RM ANOVA per mouse | day x  genotype | interaction F(2, 24)=26.22  day F(2, 24)=0.13 genotype F(1,12)=0.67 | p<0.0001  p=0.8783  p=0.4267 | Bonferroni's multiple comparisons test | YES |
| 3E | Time in the warm spot (%) | Dyn^Cre^ (n=7 mice) vs GFP (n=7mice) | unpaired t test, two tailed |  | t(12)=0.38 | p=0.70 |  | YES |
| 3F | Time in the center of OF (%) | Dyn^Cre^ (n=7 mice) vs GFP (n=7mice) | unpaired t test, two tailed |  | t(12)=2.63 | p=0.02 |  | YES |
| 3G | Distance moved in the OF (m) | Dyn^Cre^ (n=7 mice) vs GFP (n=7mice) | unpaired t test, two tailed |  | t(12)=0.9323 | p=0.3696 |  | YES |
| 3H | Time sniffing NO (%) | Dyn^Cre^ (n=7 mice) vs GFP (n=7mice) | unpaired t test, two tailed |  | t(12)=3.57 | p=0.003 |  | YES |
| 4B | O_2_ consumption | Enk^Cre^ (n=5 mice) vs GFP (n=5 mice) | Two-way RM ANOVA per mouse | complex x genotype | interaction F(5, 40)=8.12  complex F(5, 40)=79.17 genotype F(1,8)=12.78 | p=0.001  p<0.0001  p=0.007 | Bonferroni's multiple comparisons test | YES |
| 4C | ATP luminescence | Enk^Cre^ (n=5 mice) vs GFP (n=5 mice) | Mann-Whitney |  | U(11) | p=0.80 |  | NO |
| 4D | ADP luminescence | Enk^Cre^ (n=5 mice) vs GFP (n=5 mice) | Mann-Whitney |  | U(1) | p=0.01 |  | NO |
| 4E | ATP/ADP | Enk^Cre^ (n=5 mice) vs GFP (n=5 mice) | Mann-Whitney |  | U(2) | p=0.03 |  | NO |
| 4G | IP3-VDAC1 interactions (dots/nuclei) | Ctrl (n=7, Mfn2^f/f^)  vs Enk^Cre^ (n=3 mice) | Mann-Whitney |  | U(0) | p=0.0167 |  | NO |
| 5B | Number of intersections, Sholl profile | Mfn2^f/f^ (n=14) vs Mfn2^Adora2a^ (n=12) | Two-way RM ANOVA per cell | radius x genotype | interaction F(25, 624)=2.07 radius F(25, 624)=155.9 genotype F(1, 624)=106.2 | p<0.001  p<0.0001  p<0.0001 | Bonferroni's multiple comparisons test | YES |
| 5C | Total dendritic length | Mfn2^f/f^ (n=12) vs Mfn2^Adora2a^(n=14) | unpaired t test per cell, two tailed |  | t(24)=5.34 | p<0.0001 |  | YES |
| 5F | Cum. frequency of interevent-interval | GFP (N=5 mice, n=12 cells) vs Enk^Cre^ (N=6 mice, n=18 cells) | Kolmogorov-Smirnov test |  | D=0.1192 | p<0.0001 |  | YES |
| 5G | Cum. frequency of peak amplitude | GFP (N=5 mice, n=12 cells) vs Enk^Cre^ (N=6 mice, n=18 cells) | Kolmogorov-Smirnov test |  | D=0.1098 | p<0.0001 |  | YES |
| 5H | Cum. frequency of rise time | GFP (N=5 mice, n=12 cells) vs Enk^Cre^ (N=6 mice, n=18 cells) | Kolmogorov-Smirnov test |  | D=0.05443 | p=0.0007 |  | YES |
| 5J | Firing frequency | GFP (N=6 mice, n=15 cells) vs Enk^Cre^ (N=7 mice, n=17 cells) | Two-way ANOVA per cell | ramp x genotype | interaction F(5,180)=0.6235  ramp F(5,180)=19.52  genotype F(1,180)=15.58 | p=0.6820  p<0.0001  p=0.0001 |  | YES |
| 5K | Rheobase | GFP (N=6 mice, n=15 cells) vs Enk^Cre^ (N=7 mice, n=17 cells) | unpaired t test per cell, two tailed |  | t(21.16)=2.485 | p=0.0214 |  | YES |
| 5L | Firing threshold (V_thres_) | GFP (N=6 mice, n=15 cells) vs Enk^Cre^ (N=7 mice, n=17 cells) | unpaired t test per cell, two tailed |  | t(28.25)=2.25 | p=0.0324 |  | YES |
| S1B | Distance traveled in EPM (m) | LA (n=7 rats) vs HA (n=7 rats) | unpaired t test, two tailed |  | t(12)=0.0956 | p=0.9254 |  | YES |
| S1C | Time in the centre of the OF (%) | LA (n=7 rats) vs HA (n=7 rats) | unpaired t test, two tailed |  | t(12)=2.654 | p=0.021 |  | YES |
| S1D | Latency to sniffing the NO (s) | LA (n=7 rats) vs HA (n=7 rats) | unpaired t test, two tailed |  | t(12)=2.42 | p=0.0323 |  | YES |
| S1E | Social dominance (%) | LA (n=5 rats) vs HA (n=5 rats) | unpaired t test, two tailed |  | t(8)=2.352 | p=0.0464 |  | YES |
| S1F | Correlation matrix | (n=12 rats) | Pearson correlation | social domin. column | | p=0.618  p=0.002  p=0.002  p<0.001 |  | YES |
|  |  |  |  | social pref. column | | p=0.427  p=0.434  p=0.699 |  |  |
|  |  |  |  | time in open arms column | | p<0.0001  p<0.001 |  |  |
|  |  |  |  | mfn2 levels in NAc column | | p=00478 |  |  |
| S2A | Time in the tube (s) | Enk^Cre^ (n=7 mice) vs GFP (n=8 mice) | Two-way RM ANOVA per mouse | day x  genotype | interaction F(5, 56)=0.362  day F(5, 65)=4.194 genotype F(1,13)=1.457 | p=0.8724  p=0.0023  p=0.3254 |  | YES |
| S2B | Latency to fall (s) | Enk^Cre^ (n=8 mice) vs GFP (n=8 mice) | unpaired t test, two tailed |  | t(14)=0.7522 | p=0.4644 |  | YES |
| S2C | Body weight (g) | Enk^Cre^ (n=8 mice) vs GFP (n=8 mice) | unpaired t test, two tailed |  | t(14)=0.8176 | p=0.4273 |  | YES |
| S2D | Corticosterone (ng/ml) | Enk^Cre^ (n=8 mice) vs GFP (n=8 mice) | unpaired t test, two tailed |  | t(14)=0.2371 | p=0.816 |  | YES |
| S2E | Testosterone (ng/ml) | Enk^Cre^ (n=8 mice) vs GFP (n=8 mice) | unpaired t test, two tailed |  | t(14)=1.778 | p=0.0972 |  | YES |
| S3A | Time in the tube (s) | Dyn^Cre^ (n=7 mice) vs GFP (n=7 mice) | Two-way RM ANOVA per mouse | day x  genotype | interaction F(5, 60)=0.362  day F(5, 60)=10.37 genotype F(1,12)=0.6275 | p=0.6494  p<0.0001  p=0.4437 |  | YES |
| S3B | Latency to fall (s) | Dyn^Cre^ (n=7 mice) vs GFP (n=7 mice) | unpaired t test, two tailed |  | t(12)=0.2114 | p=0.8361 |  | YES |
| S3C | Body weight (g) | Dyn^Cre^ (n=7 mice) vs GFP (n=7 mice) | unpaired t test, two tailed |  | t(12)=0.4254 | p=0.6787 |  | YES |
| S3D | Corticosterone (ng/ml) | Dyn^Cre^ (n=7 mice) vs GFP (n=7 mice) | unpaired t test, two tailed |  | t(12)=0.9549 | p=0.3585 |  | YES |
| S3E | Testosterone (ng/ml) | Dyn^Cre^ (n=7 mice) vs GFP (n=7 mice) | unpaired t test, two tailed |  | t(12)=0.6014 | p=0.5588 |  | YES |
| S4D’ | NeuN+ cell/unit area | ctr (n=5 mice) vs Enk^Cre^ (n=5 mice) vs Dyn^Cre^ (n=5 mice) | One-way ANOVA |  | F(2,12)=2.619 | p=0.1138 |  | YES |
| S4E’ | NeuN+ cell/unit area | ctr(n=5 mice) vs Enk^Cre^ (n=5 mice)  ctr (n=5 mice) vs Dyn^Cre^ (n=5 mice) | one sample t test  one sample t test |  | t(4)=0.5056  t(4)=0.6370 | p=0.6396  p=0.5588 |  | YES |

**Table S1. Details on statistical analyses**
